# Kinetics of HIV-Specific CTL Responses Plays a Minimal Role in Determining HIV Escape Dynamics

**DOI:** 10.1101/194886

**Authors:** Yiding Yang, Vitaly V. Ganusov

**Affiliations:** Department of Microbiology, University of Tennessee, Knoxville, TN 37996, USA; National Institute for Mathematical and Biological Synthesis University of Tennessee, Knoxville, TN 37996, USA; Department of Mathematics, University of Tennessee, Knoxville, TN 37996, USA

**Keywords:** HIV, CTL escape, multiple responses, mathematical model, model fitting, likelihood

## Abstract

Cytotoxic T lymphocytes (CTLs) have been suggested to play an important role in controlling human immunodeficiency virus (HIV-1 or simply HIV) infection. HIV, due to its high mutation rate, can evade recognition of T cell responses variants that can not be recognized by HIV-specific CTLs. Although HIV escape from CTL responses has been well documented, factors contributing to the timing and the rate of viral escape from T cells have not been fully elucidated. Fitness costs associated with escape and magnitude of the epitope-specific T cell response are generally considered to be the key in determining timing of HIV escape. Several previous analyses generally ignored the kinetics of T cell responses in predicting viral escape by either considering constant or maximal T cell response; several studies also considered escape from different T cell responses to be independent. Here we focus our analysis on data from two patients from a recent study with relatively frequent measurements of both virus sequences and HIV-specific T cell response to determine impact of CTL kinetics on viral escape. In contrast with our expectation we found that including temporal dynamics of epitope-specific T cell response did not improve the quality of fit of different models to escape data. We also found that for well sampled escape data the estimates of the model parameters including T cell killing efficacy did not strongly depend on the underlying model for escapes: models assuming independent, sequential, or concurrent escapes from multiple CTL responses gave similar estimates for CTL killing efficacy. Interestingly, the model assuming sequential escapes (i.e., escapes occurring along a defined pathway) was unable to accurately describe data on escapes occurring rapidly within a short-time window, suggesting that some of model assumptions must be violated for such escapes. Our results thus suggest that the current sparse measurements of temporal CTL dynamics in blood bear little quantitative information to improve predictions of HIV escape kinetics. More frequent measurements using more sensitive techniques and sampling in secondary lymphoid tissues may allow to better understand whether and how CTL kinetics impacts viral escape.

**Abbreviations**
CTL: cytotoxic T lymphocyte
HIV: human immunodeficiency virus
SIV: simian immunodeficiency virus

## 1 Introduction

In 2014, the number of people living with human immunodeficiency virus 1(HIV-1 or simply HIV) was estimated as 36.9 million [50], with roughly 2 million new HIV infections and 1.2 million people dead of HIV-induced diseases (AIDS) [51].Cytotoxic CD8+ T lymphocyte (CTL) responses play an important role in control of virus replication [6, 38] by modulating some important predictors of disease progression (e.g., viral set-point and the rate of CD4+ loss rate [46]). Generation of HIV-specific CD8+ T cells by vaccination is one of the current approaches in developing HIV vaccines [23, 49]. However, HIV is able to generate mutants (termed “CTL escape mutants”) that are not recognized by HIV specific T cells, which may be one of the reasons for failure of T cell based vaccines [3, 21, 44]. Better understanding of mechanisms of viral escape and principles governing CD8+ T cell responses to HIV may allow us to evaluate *in silico* a potential efficacy of T cell based HIV vaccines.

Viral escape from CTL responses follows a somewhat predictive pattern with more dominant (larger magnitude) CTL responses leading to earlier viral escape [4, 31]. However, not every CTL response elicits an escape and sometimes viral mutations occur in regions predicted to be recognized by CTLs but in the absence of detectable response [20].To understand the timing and kinetics of CTL escape in HIV/SIV infection, mathematical models have been proposed previously on the dynamics of viral escape from a single CTL response (e.g., [2, 12, 15–17, 33, 42]). These initial models made a strong assumption of independent viral escape — i.e., it was assumed that viruses escaping from different CTL responses do not compete. Recent work, however, suggested presence of clonal interference and genetic hitchhiking among immune escape variants through reconstruction of HIV whole genome haplotypes [39], and similar concurrent CTL escapes were observed in four HIV-infected patients [29]. Clonal interference was suggested to impact the estimates of the escape rates [18, 19]. Even though several models have been developed to describe the dynamics of escapes from multiple CTL responses (e.g., [16–19, 26, 48]), many of these studies involved only model simulations and did not use information on the actual kinetics of HIV-specific CTL responses in predicting viral escape.

Here we explored whether including experimentally measured CTL kinetics improves description of the viral escape data. In doing so we compared predictions of three alternative models of viral escape from CTL responses such as independent escapes, sequential escapes, and concurrent escapes. In the first model (independent escapes) we assumed that escape from any given CTL response occurs independently of other escapes and directly from the wild-type, i.e., we ignored the effects of clonal interference – in essence assuming high effective population size and/or high recombination rate. Of note, several recent experimental papers also assumed independent escapes [4, 20, 31]. In the second model (sequential escape) we assumed that escapes from different CTL responses occur along a defined pathway, generally set by the sequences of escape occurrence in the data. This model assumes strong clonal interference which may arise at low effective population size or when recombination rate is low. Finally, in the third model (concurrent escape) we tracked all escape variants simultaneously thus allowing for co-existence of multiple escape variants (i.e., escapes could occur along multiple alternative pathways). Interestingly, we found that for well sampled data on virus evolution the estimated CTL killing efficacies were independent of the model for viral escape. Some escape data could not be well described by the sequential escape model for biologically reasonable parameters. Furthermore, explicitly taking CTL kinetics into account did not improve the quality of fit of different models to escape data. Our results suggest that CTL kinetics in the blood as it is currently available may bear limited information relevant to improve description of kinetics of HIV escape from CTL responses.

## 2 Materials and Methods

### 2.1 Experimental data

Experimental details of patient enrollment and data collection were described in detail previously [20, 31]. In short, data from 17 patients in the Center for HIV/AIDS Vaccine Immunology (CHAVI) infected acutely with HIV 1 (subtypes B or C) were analyzed in great detail. All patients were infected with a single transmitted/founder (T/F) virus as determined by the single genome amplification and sequencing (SGA/S), and there were enough samples to accurately quantify CTL response to the whole viral proteome. In each patient, the kinetics of virus specific CTL (CD8+ T cell) responses were measured using peptide stimulated IFN-γ ELISPOT assay and/or intracellular cytokine staining (ICS) six months after enrollment using peptides matched to the founder virus sequence [20, 31]. For CTL responses measured by ELISPOT, the reported magnitude of the response was the number of cells, producing IFN-γ, per 10^6^ peripheral blood mononuclear cells (PBMC). Multiple viruses were sequenced by SGA/S, and all sequences were compared at cites coding for CTL epitopes and changes in the percentage of transmitted (wild-type) sequences were followed over time [31]. The dynamics of the HIV specific CTL responses and viral escape from epitope specific CTL responses were measured longitudinally. Escape mutants were identified as viral variants with mutations in regions recognized by patient‚s CTL responses with a reduced (or fully abrogated) production of IFN-γ following T cell stimulation. In many cases mutation in a single position was responsible for the escape. In our analysis all viral variants which did not have the wild-type amino acid in the epitope region were considered as escape variants.

Review of the virus evolution and CTL dynamics data in all 17 patients revealed some data limitations. In particular, data for many patients lacked adequate temporal resolution to accurately estimate virus escape rates. In the vast majority of viral escape variants, escapes often occurred rapidly between two sequential time points with the frequency of the escape variant jumping from 0 to 1. While previously it was suggested that such data may be modified to provide an estimate of the escape rate [2, 12, 16], such approaches may lead to biased parameter estimates [26]. While development of a method for unbiased estimation of escape rate from sparse data is ongoing (Ganusov et al., ms. in preparation), for this analysis we focused on patients CH131 and CH159 in which viral escape rates could potentially be accurately estimated due to sufficiently frequent sampling. While data from these patients were presented before [31], linking of escape and CTL response dynamics was not yet performed.

### 2.2 Model of viral escape from a single CTL response

Models describing the dynamics of viral escape from a single cytotoxic T lymphocyte (CTL) response have been developed and adopted by different researchers (e.g., [2, 12, 15–17]). Here we start with the basic model formulated earlier [17], and extend it to viral escape dynamics from multiple CTL responses. The model of viral escape from a single CTL response can be extended from the basic viral dynamics model [40] in the following way:

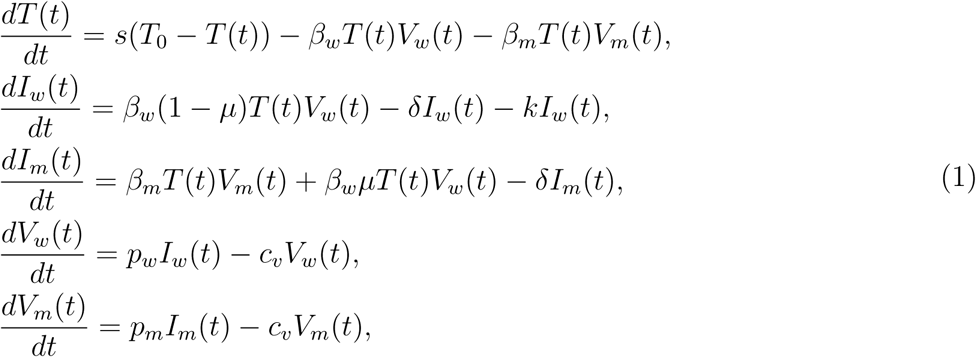

where *T* (*t*) is the density of uninfected target cells; *I*_*w*_(*t*) and *I*_*m*_(*t*) is the density of target cells infected by the wild-type or escape variant viruses, respectively; *V*_*w*_(*t*) and *V*_*m*_(*t*) is the density of wild-type or escape variant viruses, respectively; *s* is the turnover rate of uninfected target cells; *T*_0_ is the preinfection level of uninfected target cells; *β_w_* and *β_m_* is infection rate of wild-type or escape variant viruses, respectively; *µ* is the probability of mutation from wild-type to escape mutant during reverse transcription of viral RNA into proviral DNA; *δ* is the death rate of infected cells due to viral pathogenicity; *k* is the killing rate of wild-type virus infected cell due to CTL response; *p*_*w*_ and *p*_*m*_ is the rate at which cells infected by wild-type or escape mutant viruses produce viruses; and *c*_*v*_ is the clearance rate of free viral particles.

In this model (eqn. (1)), we assume that target cells infected by wild-type (*V*_*w*_(*t*)) and escape viruses (*V*_*m*_(*t*)) differ by two factors: viral infectivity (*β_w_* and *β_m_*) and the rate of virus production (*p*_*w*_ and *p*_*m*_). Given that *in vivo* viral particles are short-lived [41, 43], to a good approximation we may assume a quasi steady state for the virus particle concentration leading to 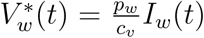 and 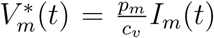. We define a fitness cost 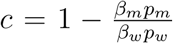 where *c* can be positive or negative. Positive *c* means true fitness cost of escape mutations, that is escape variant has a lower replication rate (*β_m_p_m_ ≤β_w_p_w_*) [45], and negative *c* implies fitness advantage of escape virus [45, 52]. By straightforward calculation, the system (eqn. (1)) can be written as

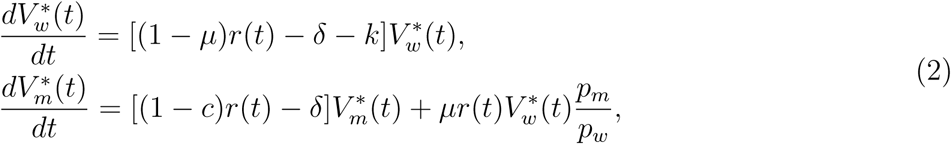

For convenience, we replace 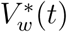 and 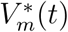by *w*(*t*) or *m*(*t*), respectively, and assume that the wild-type and escape viruses differ only in the rate of infectivity (that is *β_w_* ≥ *β_m_* and *p*_*w*_ = *p*_*m*_) [20], the system (2) can be simplified as

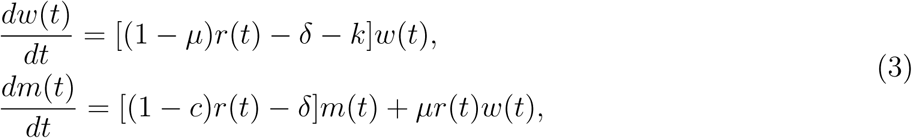

where 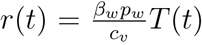 is the replication rate of cells infected by wild-type virus, and 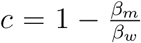 is the cost of the escape mutation defined as a selection coefficient. The frequency of the escape variant in the whole population is given by 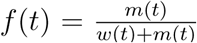. This is perhaps the simplest model for a viral escape from a single CTL response. This is denoted as **model 1** in the paper.

### 2.3 Models of viral escapes from multiple CTL responses

Mathematical model given in eqn. (3) tracks changes in densities of wild-type virus and a single variant that has escaped recognition from a single epitope specific CTL response. In acute HIV infection, the virus can escape from recognition of multiple CTL responses, which are specific to several viral epitopes [20, 47]. Several models have been developed to describe the dynamics of escapes from multiple CTL responses (e.g., [16, 17, 48]). Our model is an extension of previous models [16, 17] incorporating mutations from wild-type virus to different viral escapes. In contrast with previous studies in our analyses here we used experimentally measured time courses of different CTL responses [31].

To track the dynamics of viral escape from multiple responses, we assume that there are in total *n* CTL responses that control viral growth, and virus can potentially escape from all *n* responses. We use *m***_i_** to denote the density of variants where **i** is a vector **i** = (*i*_1_, *i*_2_, *…, i*_*n*_) denoting the positions of *n* epitopes, and we define *i*_*j*_ = 0 if there is no mutation in the *j*^*th*^ CTL epitope and *i*_*j*_ = 1 if there is a mutation leading to an escape from the *j*^*th*^ (1 ≤ *j* ≤ *n*) CTL response. We denote the set of escape variant as *I*, that is **i** *∈ I*. The wild-type variant is then denoted as (0, 0, *…* 0).

For our analysis, we neglect recombination and backward mutation from mutant to wild-type. We use *k*_*i*_, *c*_*i*_ and µ_*i*_ to denote killing rate due to *i*^th^ CTL response, cost of escape mutation from the *i*^th^ CTL response and mutation rate for the *i*^th^ epitope, respectively. Due to a small rate of double mutation [34], we assume that escape virus is generated with only one mutation in a single generation. That is for two escape variants *m***_i_** = *m*_(*i*_1_,*i*_2_,…,*i_n_*)_ and *m***_j_** = *m*_(*j*_1_,*j*_2_,…,*j*_n_)_, we define the mutation rate *M***_i,j_** from *m***_i_** to *m***_j_** as *µ_k_*, if and only if *m***_j_** has only one more mutation at position *k* than *m***_i_** and all other positions are exactly same. For example, when there are 3 CTL responses, the mutation rate from *m*_(1,0,0)_ to *m*_(1,1,0)_ is *µ*_2_, and the mutation rate from *m*_(0,0,0)_ to *m*_(1,0,1)_ is 0. Assuming multiplicative fitness (detailed deviation is given in Section S2 in Supplement), that is, the fitness cost of a variant **i** = (*i*_1_, *i*_2_, *…, i*_*n*_) is 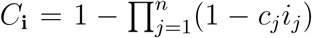. The death rate of the escape variant **i** = (*i*_1_, *i*_2_, *…, i*_*n*_) due to remaining CTL responses is given by 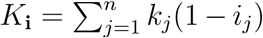 where we assume that killing of infected cells by different CTL responses is additive.

Similar to eqn. (3), the dynamics of the wild-type and escape variants are given by

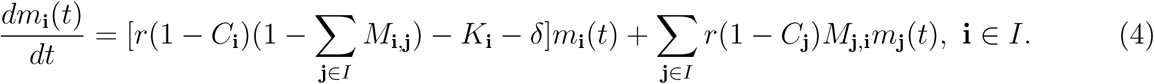

We define *M* (*t*) = ∑_**i**∈*I*_ *m***_i_** as the total density of all variants in the opulation, and *f*_*j*_(*t*) (*j* = 1, *…, n*) is the fraction of viral variants that have escaped recognition from the *j*^*th*^ CTL response.The frequency of a viral variant escaping from the *j*^*th*^ response is given by

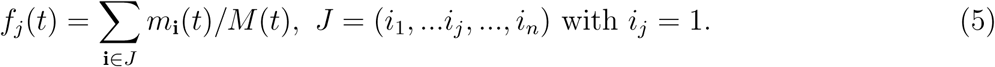

Based on previous work [26, 28, 29], we assume that there are two alternative ways to generate escape mutants (Figure 1). The first way can be called “sequential” escape (**model 2**), that is escape mutants are generated sequentially along a defined path from wild-type viruses. This is likely to happen when the effective population size of HIV is small and when the rate of recombination is negligible. The second way can be described as “concurrent” escape (**model 3**), in which the virus can escape from *n* CTL responses simultaneously along multiple different pathways. This is likely to happen when the HIV effective population size is large. With *n* CTL responses, there are *n* escape Variants for “sequential” escape and 2^*n*^ *-*1 escape variants for “concurrent” escape in addition to the wild-type variant. For example, with *n* = 3 CTL responses, for “sequential” escape there are 3 escape variants: *m*_(1,0,0)_, *m*_(1,1,0)_, and *m*_(1,1,1)_ with *m*_(0,0,0)_ being the wild-type virus. For “concurrent” escape there are 7 escape variants: *m*(1,0,0), *m*(0,1,0), *m*(0,0,1), *m*(1,1,0), *m*(1,0,1), *m*(0,1,1) and *m*(1,1,1) with *m*_(0,0,0)_ being the wild-type virus (Figure 1). Detailed equations for both models with *n* = 3 CTL responses can be found in Supplement (Section S2). It is interesting to note that > escape is a simplification of “concurrent” escape when the effective population size is small. Previous work did not fully resolve whether CTL escapes in HIV infection occur sequentially of concurrently [26, 29]; most likely the type of escape varies by patient.

**Figure 1:**
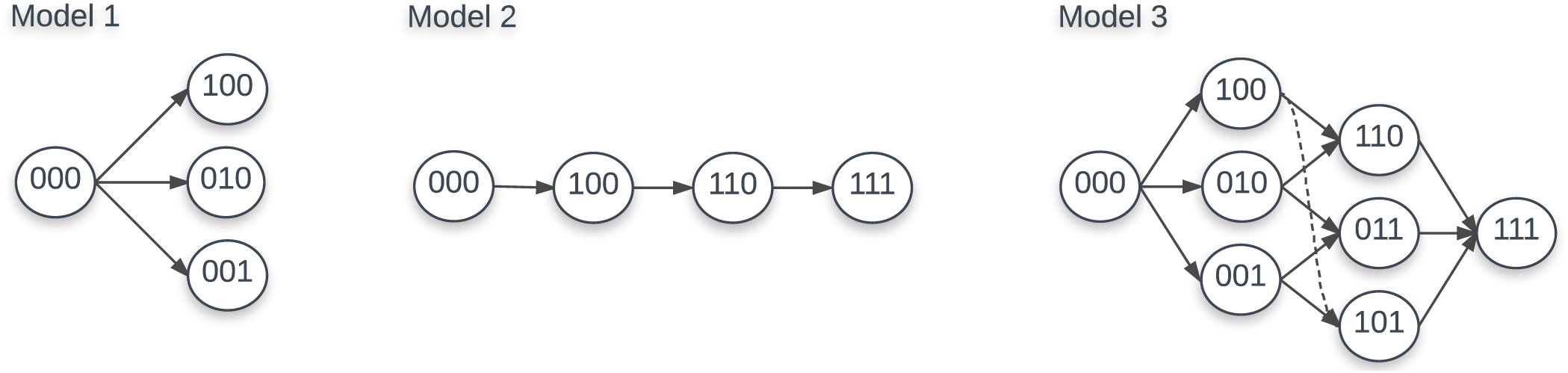
Escape paths for models 1, 2 & 3 with 3 CTL responses. For model 1, there are 3 escape variants:*m*(1,0,0), *m*(0,1,0) and *m*(0,0,1).For model 2 there are also 3 escape variants:*m*(1,0,0), *m*(1,1,0), and *m*(1,1,1).For model 3 there are 7 escape variants: *m*(1,0,0), *m*(0,1,0), *m*(0,0,1), *m*(1,1,0), *m*(1,0,1), *m*(0,1,1) and *m*(1,1,1).In each case, *m*(0,0,0) is the wild-type virus.

## Models for CTL response

The killing rate *k*_*i*_ of the CTL response specific to the *i*^*th*^ epitope in all three models is composed of two parts: the per cell killing efficacy of CTLs 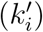 and the number of epitope-specific CTLs (*E*_*i*_) [15]. Previously the killing rates *k*_*i*_ were often set to a constant (e.g., [15, 17]), or were set to a certain form 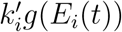 where *g*_*i*_(*E*_i_*t*)) is a function of epitope specific CTL responses *E*_*i*_(*t*) (e.g., [1, 19]). With the measured epitope-specific CTL response dynamics [20], we adopted two forms of killing rate: constant *k*_*i*_ (termed as “constant response”) or time-dependent killinrate 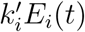 (termed as “interpolated/fitted response”). We used the “mass action” killing term to describe effect of CTLs on virus dynamics because it is the simplest form, it involves minimum parameters, and it is supported by some experimental data [14].

Based on the available time course information of epitope-specific T cell response *E*_*i*_(*t*), we used the first-order interpolation function (termed as “interpolated response”) or the fitted response function (termed as “fitted response”) by the *T*_on_ *T*_off_ model [10] to quantify the kinetics of HIV-specific CTL responses. The *T*_on_ *T*_off_ model assumes that the response starts with *E*_0_ epitope specific CD8+ T cells that become activated at time *T*_on_. Activated T cells start proliferating at a rate *ρ* and reach the peak at time *T*_off_. After the peak, epitopes specific CD8+ T cells decline at a rate *α*.The dynamics of the CD8+ T cell response *E*(*t*) is given thus by the following differential equation:

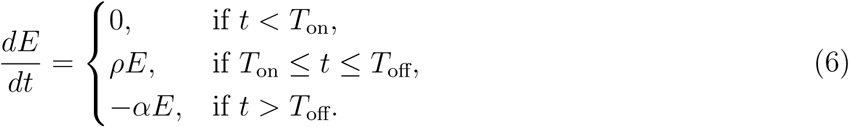

with *E*(0) = *E*_0_. Here the “precursor frequency” *E*_0_ is a generalized recruitment parameter, which combines the true precursor frequency and the recruitment rate/time [9, 10]. Our recent work showed that this model (eqn. (6)) reasonably well describes kinetics of HIV-specific CTL responses in acute HIV infection (Yang and Ganusov (in review)). When fitting the model (eqn. (6)) to experimental data of CTL dynamics we changed all initial undetected response values from 0 to 1; the latter was the detection limit in the data.

### 2.5 Statistics

Previously, under the assumption that some mutants are present initially, researchers (e.g., [1, 15]) fit a logistic model to data on viral escape kinetics by the method of nonlinear least squares [5]. In essence, this is a maximum likelihood method which assumes normally distributed residuals. While this standard statistical method provides reasonable parameter estimates it assumes equal weights to different data points independently of how many viral sequences were measured at every time point which is likely to be unrealistic for most experimental studies. Here we follow the method proposed recently [17] to use binomial distribution (and thus different weights for different measurements/time points) in the likelihood of the model given the escape data. For HIV escape from a single CTL response the log-likelihood function is given by

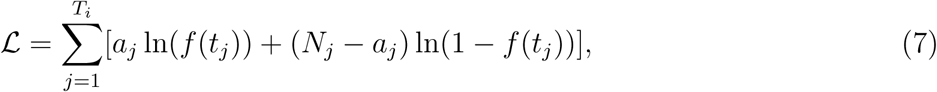

where *a*_*j*_ is the number of escape variant sequences in a sample of *N*_*j*_ sequences at the sample time *t*_*j*_, *T*_*j*_ is the number of measured time points for a specific viral escape trajectory, and *f* (*t*_*j*_) is the predicted frequency of a specific viral escape variant at time *t*_*j*_. Model parameters were thus found by maximizing the log-likelihood function (eqn. (7)).

To discriminate between alternative models under different parameter constrains we used corrected Akaike information criterion (AIC) scores [8]. The model fit with the minimum AIC score among tested models was treated as the best model; however, a difference of less than 3 AIC units is generally viewed as not significant [8]. To test the statistical significance of the differences between parameters found by fitting different models, we used a bootstrap approach [11]. In this approach we resampled the data 1000 times using the Random routine in Mathematica assuming beta distribution for sequencing data [7], fitted models to bootstrap samples, and recorded all estimated parameters. For the same parameter, we use either paired and unpaired t-test to compare the means from different models.

Both fitness costs of escape mutations and the killing efficacy of the CTL response determine the kinetics of viral escape from T cells [2, 12, 15], and that viral escape (sequence) data in most cases are not sufficient to estimate both rates [15]. Therefore, in our analyses to avoid overfitting we set fitness cost of escape to zero *c*_*i*_ = 0. In all fits we assumed that the rate of virus replication *r* = 1.5/day [40].

While multiple models may be able to describe accurately experimental data, some models may do so at biologically unreasonable parameters. For example, estimated rate of mutation at different epitopes may be unrealistically large. Thus, in our analysis we assume that mutation rates which are above 10^-3^ are likely to be unrealistic given that currently estimated HIV mutation rate is about 3.2×10^-5^ per bp per generation [34] and size of a CTL epitope is 8-10 amino acids (3×10×3.2×10^-5^ ≈10^-3^).

To fit the *T*_on_-*T*_off_ model (eqn. (6)) to experimental data using non-linear least squares we log-transformed the model predictions and the data.

When interpolating CTL response kinetics, there was often not enough information on the starting point (day 0). In such situations we set the initial CTL density as 1 (the detection level for this data set) for simplicity. Other starting points (e.g., intersection point of the CTL response axis and the reverse extension line of the interpolation function) were also tested and led to similar results (not shown). This was largely due to the fact that in our models CTLs at low densities are not expected to exert large selective pressure on the virus population due to assumed mass action killing term.

## 3 Results

### 3.1 Statistical model impacts estimation of the escape (killing) rate

Given virus evolution data we may be often interested in quantifying selecting pressures driving specific changes in the virus population. Following HIV -1 infection, the virus escapes from several cytotoxic T lymphocyte (CTL) responses [36], and multiple studies used mathematical models of various levels of complexity to estimate the predicted efficacy at which CTLs recognize and eliminate cells, infected with the wild-type (unescaped) virus [2, 12, 15–17, 26]. Many of these previous studies estimated the rate of HIV escape from immunity using nonlinear least squares which explicitly assumes normal distribution of the deviations between model predictions and data [2, 12, 15, 16]. However, the assumption of normally distributed residuals is likely to be violated for data when only a handful of viral genomes are sequences – which is common in many studies involving single genome amplification and sequencing techniques (SGA/S). We have recently proposed to use a likelihood approach which assumes virus genome sampling to follow a binomial distribution [17]. This binomial distribution-based likelihood approach showed to impact the estimates of the CTL killing rate (escape rate can be proportional to the killing rate under an assumption of constant CTL response) when compared to normal distribution based likelihood approach (least squares) [17]. However, this previous comparison was done on data which were fairly sparse and comparison involved modifications of data to allow for non-zero and non-one frequencies of the escape variant [2, 12], and thus, it remained unclear if estimates of escape rates are truly dependent on the statistical model for better sampled data.

Unfortunately, in our cohort of 17 patients [31] very few patients were sampled frequently enough to observe gradual accumulation of escape variants in the population (i.e., data with two sequential time points with mutant frequency in the range 0 < *f* < 1 were rare). For the analysis we, therefore, used the escape data from two patients, CH131 and CH159, where CTL and HIV sequence measurements were sufficiently frequent to address our modeling questions. We fitted a simple mathematical model describing escape of the virus from a single constant (non-changing) CTL response (eqn. (3)) to the data from one patient CH159 (Figure 2) assuming two different statistical models: with normally distributed residuals (least squares) or binomial distribution based likelihood (eqn. (7)). Consistent with our previous observation we found that the type of statistical model impacts the estimate of the escape rate (*k* in Figure 2) with difference being nearly 2 fold (*k* = 0.27/day vs. *k* = 0.51/day). It is interesting to note that visually, the least squares method appear to describe the data better by accurately fitting the points with intermediate frequency of the escape variant in 20 30 days after the symptoms (but missing the another intermediate data point (12, 0.08)). However, this visually better fit is not supported by the statistics: likelihood of the model for these data is -12.64 or -10.53 for normal (Figure 2A) or binomial (Figure 2B) distribution, respectively (and AIC scores being 31.0 vs. 26.8, respectively). Interestingly, the main difference in the estimated escape rates was driven by just one data point ((*t, f*) = (12, 0.08)); removing this data point from the data led to identical estimates of the escape rate, *k* = 0.51/day, from two statistical models (results not shown). This is not surprising because with this data point removed, the information on escape rate is only coming from two data points when the frequency of the escape variant is intermediate (0 < *f* < 1).

**Figure 2:**
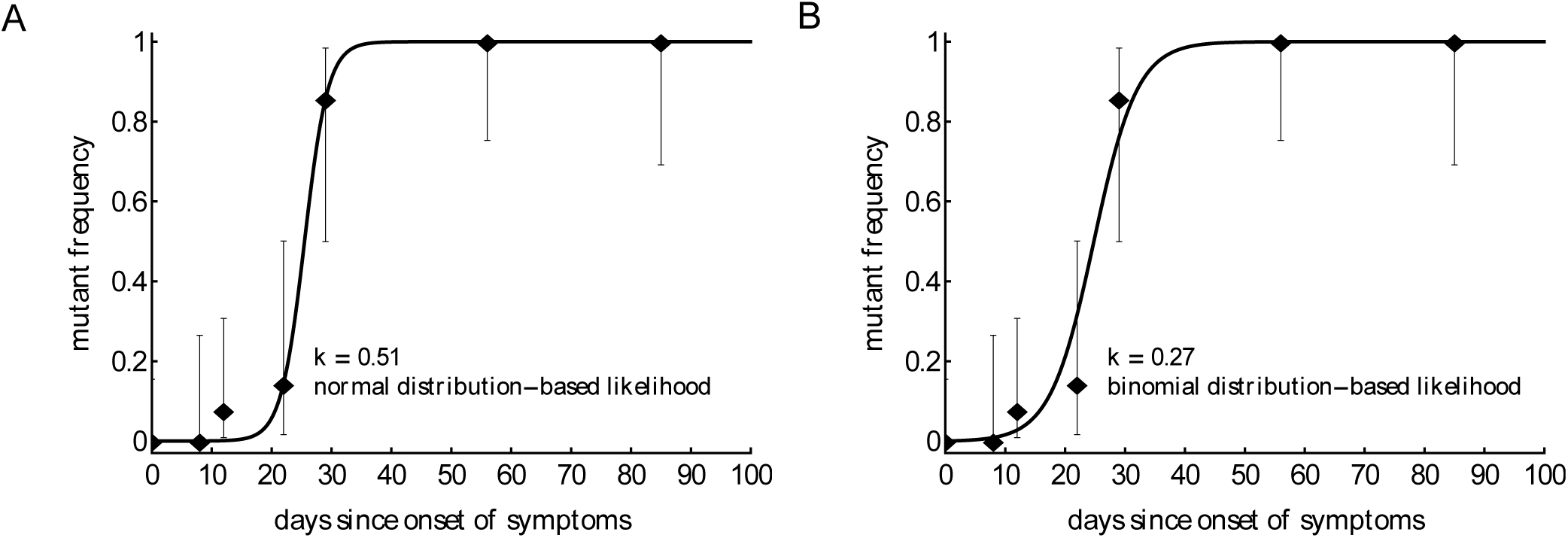
Statistical model has a strong impact on the estimated killing rate.We fit model in eqn. (7) to the same data for HIV escape in the protein region DREVLIWKFDSSLARRHL of Nef (Nef 177-194) in patient CH159, assuming normal distribution based likelihood (normally distributed residuals or nonlinear least squares, panel A) or binomial distribution based likelihood method (panel B). Data are shown as dots and bars represent the 95% confidence intervals calculated using beta distribution (Jefferey‚s intervals, [7]). The fitted parameters are *µ* = 7.76 × 10^-7^ and *k* = 0.51 day^-1^ (A), or *µ* = 2.00 × 10^-4^ and *k* = 0.27 day^-1^ (B).

As discussed before least squares may not allow to estimate escape rates, e.g. in cases when mutant frequency jumps from 0 to 1 between two subsequent time points unless data are modified [2, 12]. Similarly, models assuming normally distributed residuals may not be able to fit other types of data, in which frequency of the mutant has an intermediate value (0 < *f* < 1) at one time point only. In particular, in our analysis of another escape in patient CH159 (Rev GRPTEPVPFQLPPLERLC, see Figure 3) we could not obtain finite estimates of the escape rate using normally distributed residuals (results not shown). Rather, the model fits tended to describe accurately two data points (*t* = 22 days and *t* = 29 days) and ignore another data point (*t* = 56 days) leading to extremely high predicted escape rates (results not shown). Interestingly, using binomial distribution based likelihood allowed for an accurate fit of the model to data and the fit compromised between describing early and late data points (Figure 4A). The reason for the compromise is that a fit predicting fast escape and nearly 100% escape variant by 56 days since symptoms is highly disfavored by the binomial distribution-based likelihood because some wild-type variants were still present at day 56 (thus the weight for missing this point by the model fit was very high in binomial distribution-based likelihood but not in the normal distribution-based likelihood). Taken together, these results suggest that the type of the statistical model used to estimate HIV escape rates influences the final estimates. Therefore, many previous studies on HIV escape assuming normally distributed residuals may need to be re-evaluated for the robustness of their conclusions.

**Figure 3:**
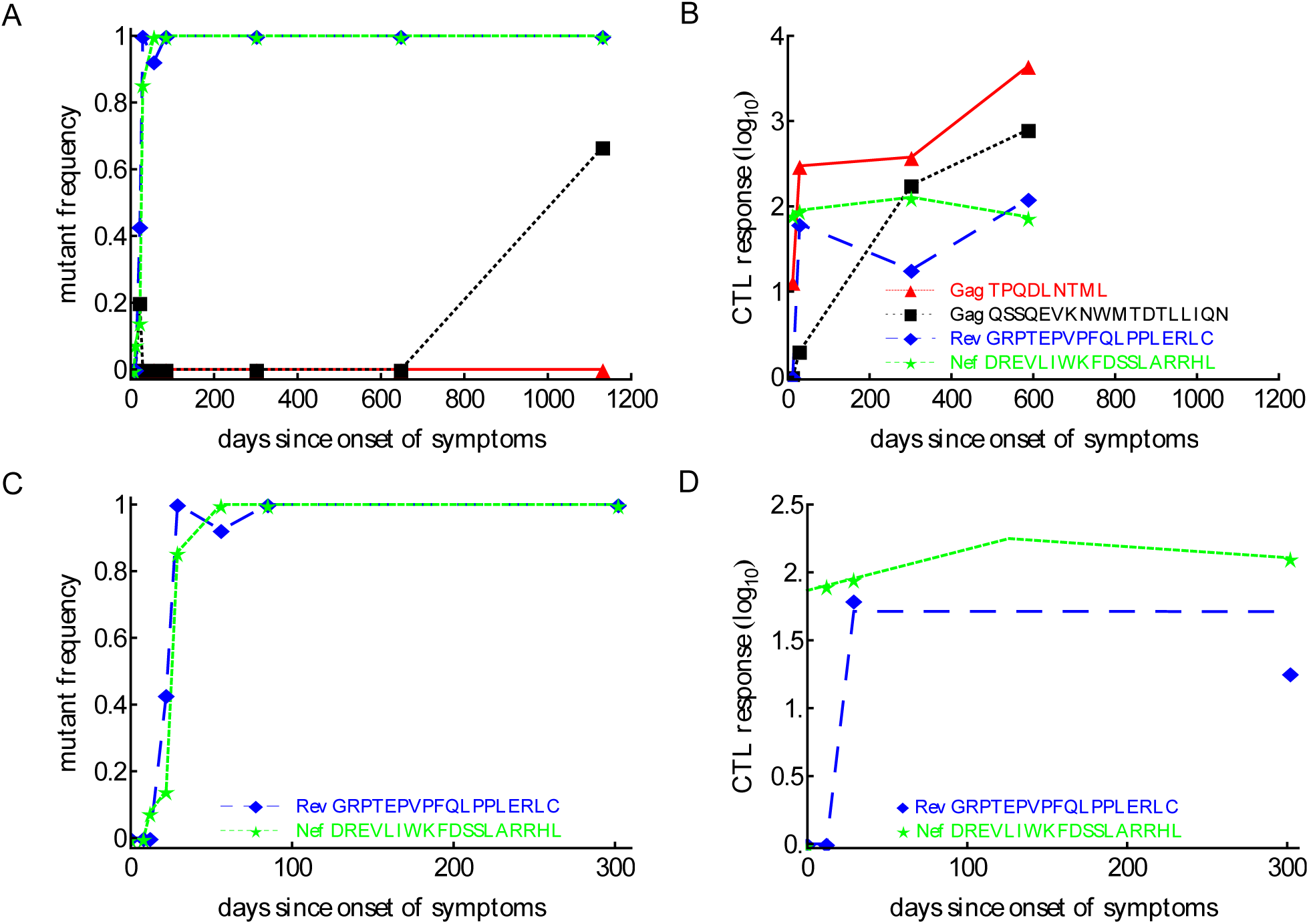
Basic dynamics of CTL response and HIV escape for patient CH159. Data are from a previous publication [31]; the data show four CTL responses in the patient (panel B) and frequencies of corresponding escape variants (panel A). Based on the selection criteria described in the Materials and in Methods we focused our analysis on CTL dynamics and escape in two regions: Rev GRPTEPVPFQLPPLERLC (65-82) and Nef DREVLIWKFDSSLARRHL (177 -194) shown for the first 200 days in panels C-D. Dashed lines in panel D are the prediction of the *T*_on_ *T*_off_ model to these data with the following estimated parameters for the Rev-specific T cell response: *E*_0_ = 1 IFN*γ*+ SFC*/*10^6^ PBMC, *T*_on_ = 12 day, *T*_off_ = 29 day, *ρ* = 0.23 day^-1^, *α* = 1.67 × 10^-6^ day^-1^; and for the Nef specific T cell response: *E*_0_ = 73.59 IFN*γ*+ SFC*/*10^6^ PBMC, *T*_on_ = 0 day, *T*_off_ = 126.05 day, *ρ* = 6.98 × 10^-3^ day^-1^, *α* = 1.86 × 10^-3^ day^-1^

**Figure 4:**
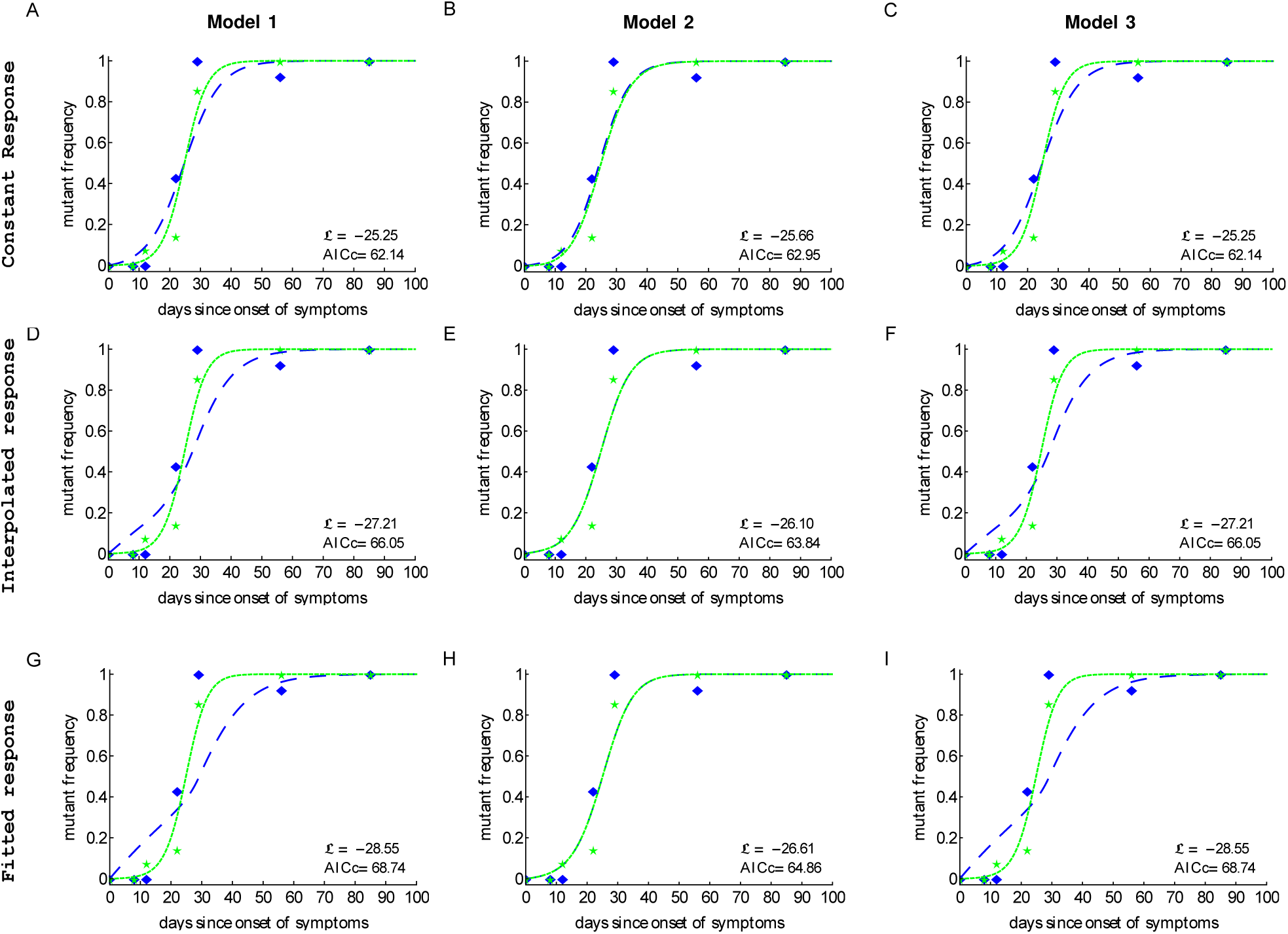
Including CTL response dynamics worsened model fits of HIV escape data in patient CH159.We fitted model 1 (independent escapes, eqn. (3)), model 2 (sequential escape, eqn. (S6)) and model 3 (concurrent escape, eqn. (S8)) to escape data in patient CH159 with different response inputs (constant, interpolated, or fitted response, see Materials and Methods for more detail). Adding direct time-dependent response (interpolated or fitted response) did not improve the quality of the model fit to data (see Table 1 for parameter estimates). Model 2 was not able to accurately describe these data for biologically reasonable mutation rates (see Table 1).

### 3.2 CTL response kinetics do not improve description of the escape data

As CTL responses drive HIV escape from epitope specific T cells, it is expected that the magnitude of the CTL response should naturally impact escape kinetics. Previous studies provided some evidence that the relative magnitude of a given CTL response in the total HIV-specific CTL response early in infection (% immunodominance) predicts the timing of viral escape [4, 31]. Immune response was also shown to impact escape of simian immunodeficiency virus (SIV) from T cell responses [32, 33, 35]. Immune response magnitude, and as a consequence, the overall CTL killing efficacy is important in determining both timing and speed of viral escape with the rate of viral escape being directly related to the immune response efficacy [15, 16]. In contrast, both initial mutant frequency, virus mutation rate, and CTL killing efficacy determine timing of viral escape [16]. Whether inclusion of the experimentally measured CTL dynamics impacts ability of mathematical models to accurately describe viral escape data has not been tested.

**Table 1:** Parameters for the three models fitted to escape data from patient CH159. Fits of the model to data are shown in Figure 4. *L*and AICc are the log-likelihood and the corrected Akaike informationcriterion value, respectively.In bold we show maximum *L* and minimum AICc reached by the models 1&3 with constant response. There are some unrealistic mutation rates given by model 2 (much bigger than10^-3^, highlighted as italic), and models 1&3 also led to slightly unrealistic mutation rates at the peptide Rev 65 82 (slightly bigger than 10^-3^). Units for *k*_*i*_ and 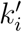 are day^-1^ and µ_*i*_ is dimensionless (same for all tables below).

|  | peptide | model 1 |  | model 2 |  | model 3 |  |
| --- | --- | --- | --- | --- | --- | --- | --- |
| constant<br>response | | mutation rate<br>($\mu_i$, i=1,2) | killing rate<br>($k_i$, i=1,2) | mutation rate<br>($\mu_i$, i=1,2) | killing rate<br>($k_i$, i=1,2) | mutation rate<br>($\mu_i$, i=1,2) | killing rate<br>($k_i$, i=1,2) |
| | Rev 65-82 | $1.68 \times 10^{-3}$ | 0.17 | $9.71 \times 10^{-4}$ | 0.20 | $1.68 \times 10^{-3}$ | 0.17 |
| | Nef 177-194 | $2.02 \times 10^{-4}$ | 0.27 | *0.11* | $6.29 \times 10^{-12}$ | $2.0 \times 10^{-4}$ | 0.27 |
| | | $\mathcal{L} = -25.25$, AICc= 62.14 | | $\mathcal{L} = -25.66$, AICc= 62.95 | | $\mathcal{L} = -25.25$, AICc= 62.14 | |
| interpolated<br>response | | mutation rate<br>($\mu_i$, i=1,2) | killing rate<br>($k'_i$, i=1,2) | mutation rate<br>($\mu_i$, i=1,2) | killing rate<br>($k'_i$, i=1,2) | mutation rate<br>($\mu_i$, i=1,2) | killing rate<br>($k'_i$, i=1,2) |
| | Rev 65-82 | $8.88 \times 10^{-3}$ | $2.12 \times 10^{-3}$ | $1.64 \times 10^{-3}$ | $2.03 \times 10^{-10}$ | $8.88 \times 10^{-3}$ | $2.12 \times 10^{-3}$ |
| | Nef 177-194 | $4.94 \times 10^{-4}$ | $3.23 \times 10^{-3}$ | *697.77* | $2.32 \times 10^{-3}$ | $4.93 \times 10^{-4}$ | $3.23 \times 10^{-3}$ |
| | | $\mathcal{L} = -27.21$, AICc= 66.05 | | $\mathcal{L} = -26.10$, AICc= 63.84 | | $\mathcal{L} = -27.21$, AICc= 66.05 | |
| fitted<br>response | | mutation rate<br>($\mu_i$, i=1,2) | killing rate<br>($k'_i$, i=1,2) | mutation rate<br>($\mu_i$, i=1,2) | killing rate<br>($k'_i$, i=1,2) | mutation rate<br>($\mu_i$, i=1,2) | killing rate<br>($k'_i$, i=1,2) |
| | Rev 65-82 | $1.43 \times 10^{-2}$ | $1.39 \times 10^{-3}$ | $1.13 \times 10^{-3}$ | $8.50 \times 10^{-18}$ | $1.43 \times 10^{-2}$ | $1.39 \times 10^{-3}$ |
| | Nef 177-194 | $2.46 \times 10^{-4}$ | $3.25 \times 10^{-3}$ | *13004.84* | $2.29 \times 10^{-3}$ | $2.47 \times 10^{-4}$ | $3.25 \times 10^{-3}$ |
| | | $\mathcal{L} = -29.68$, AICc= 70.99 | | $\mathcal{L} = -26.61$, AICc= 64.86 | | $\mathcal{L} = -29.68$, AICc= 70.99 | |

To test the benefits of using longitudinally measured CTL responses in describing viral escape data we considered several alternative models for the CTL dynamics and viral escape. Our model 1 describes the dynamics of viral escape from each CTL response independently. Models 2 & 3 describe escape from multiple CTL response that occurs sequentially or concurrently, respectively (see Materials and Methods for more details). CTL dynamics was either considered to be unimportant (i.e., killing rate *k*_*i*_ was set constant over time), or when killing rate was proportional to the experimentally measured CTL frequency (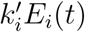), respectively. To describe CTL dynamics we either used the first order interpolation function or the *T*_on_-*T*_off_ model (eqn. (6) and see Materials and Methods for more detail).

In patient CH159, four CTL responses were detected (Figure 3B) and three of these responses were escaped within nearly 4 years of infection. Interestingly, the response specific to Gag TPQDLNTML was dominant (Figure 3B), but the corresponding escape mutant Gag TPQDLNTMLNTVGGHQAA did not appear up to 1132 days since onset of symptoms (Figure 3A).

Patient CH159 had two escape mutants in regions Rev GRPTEPVPFQLPPLERLC (Rev 65 82) and Nef DREVLIWKFDSSLARRHL (Nef 177 194) satisfying our selection criteria (Figure 3C). Despite a relative small magnitude of CTL responses specific to Rev65 and Nef177 early in infection (up to 29 days since onset of symptoms), escape mutants appeared early and their frequencies arose rapidly.

We fitted three alternative mathematical models for viral escape and three alternative models for the CTL dynamics to the data on viral escape (Figure 3C) using binomial distribution-based likelihood method (see Materials and Methods for more detail). Surprisingly, we found that the models 1 & 3 with a constant immune response described the data with best quality as judged by the AIC (or likelihood). Parameter estimates in the model 1 which assumes independent escape were nearly identical to the parameters in the model 3 which assumed concurrent escape (Figure 4 and Table 1). Importantly, adding experimentally measured CTL response dynamics (as interpolated function or by using parameterized *T*_on_ *-T*_off_ model) did not improve the quality of the model fit to escape data (Table 1). Even worse, for models 1 & 3 the fits with a fitted response were of lower quality as judged by the large increase in AIC (Table 1). Models that included an interpolated CTL response provided better fits than models with a fitted response (Table 1).

The exact reasons of why including experimentally measured CTL response dynamics led to worse fits of the escape data are unclear but perhaps rapid change in magnitude of CTL responses in this patient – if response directly impacts killing of infected cells – was simply not reflected in the kinetics of viral escape (Figure 4D&G). Specifically, CTL kinetics driven escape would predict non monotonic rise in the escape variant frequency which was not observed in the data, thus, favoring a model with a constant killing rate by CTLs.

Interestingly, the model 2 fits of the data resulted in unphysiologically large estimates for the mutation rate *µ*_2_ (Table 1). As we elaborate later (see below) this failure of the model to describe these data stems from the fact that escapes in the data occur nearly at the same time and assuming that escapes are sequential led to an unrealistic mutation rate in the second epitope. This suggests that the observed dynamics of viral escape in patient CH159 is not consistent with sequential escape.

Models 1 & 3 also predicted slightly higher than expected mutation rate *µ*_1_ (bigger than 10^-3^) for the peptide Rev 65-82. Constraining this parameter to remain *µ*_1_ ≤ 10^-3^ led to fits of significantly lower quality (likelihood ratio test, *p* < 0.05). Due to large length of the peptide, the overall mutation rate in this region could indeed be slightly higher than our calculated high bound for the mutation rate (see Materials and Methods for more detail). Furthermore, since peptide Rev 65-82 is the epitope in which first escape occurred, it was possible that the high estimate of the mutation rate could be due to late sampling of viral sequences. In these data sampling was done after patients were diagnosed with infection, however, viral escape could have started earlier and for escapes starting earlier it may be possible to describe the data with a lower mutation rate [17, 27].

Therefore, to test whether the timing of the start of the escape influences the estimate of the mutation rate we did the following. We shifted the data for two escapes forward by adding some initial zeroes to data and reverse extended the predicted CTL response curves. Then we refitted model 1 & 3 to the data under the constrain µ≤10^-3^. We found shifting the data did not improve the quality of the model fits as compared to unmodified data when CTL dynamics is explicitly taken into account as interpolated or fitted response (results not shown). However, assuming a constant response allowed to obtain lower, more physiological estimates of the mutation rate. These results suggest that inability of the models which explicitly incorporate CTL dynamics to explain kinetics of first escape with physiologically reasonable mutation rate is due to late appearance of the CTL response. Indeed, escape can only accumulate when CTL response is present and extending the time window for virus evolution but not having CTL response active will not significantly impact estimates of the mutation rate.

Given our results for one patient we next sought to investigate whether our conclusions will remain robust when looking at data from another patient. Patient CH131 had 6 CTL responses and there was escape from at least 5 of these responses in 2 years since symptoms (Figure 5). One escape, Nef EEVGFPVKPQV (Nef 64-74), occurred very early in infection, and two escapes, Env RQGYSPLS-FQTLIPNPRG (Env 709-726) and Gag VKVIEEKAFSPEVIPMFT (Gag 156-173), occurred late (Figure 5). In this patient the pattern of escape followed the ranking of immunodominance of CTL responses [31]: Nef64-specific CTLs were dominant at symptoms and drove earlier escape, while Env 709 and Gag156-specific CTLs arose later with escapes occurring later in infection (Figure 5A&B). However, there were apparently discrepancies such as two escapes in Tat epitopes (Tat DPWNH-PGSQPKTACNNCY, that is Tat 9-26 and Tat FQKKGLGISY, that is Tat 38-47) occurred at the same time while CTL responses specific to these different epitopes were of different sizes (Figure 5A&B). Because escapes in these two Tat epitopes occurred rapidly and did not have two intermediate measurements of the mutant frequency, our following analysis was only restricted to escapes in three CTL epitopes: Nef64, Env709, Gag156 (Figure 5C&D).

**Figure 5:**
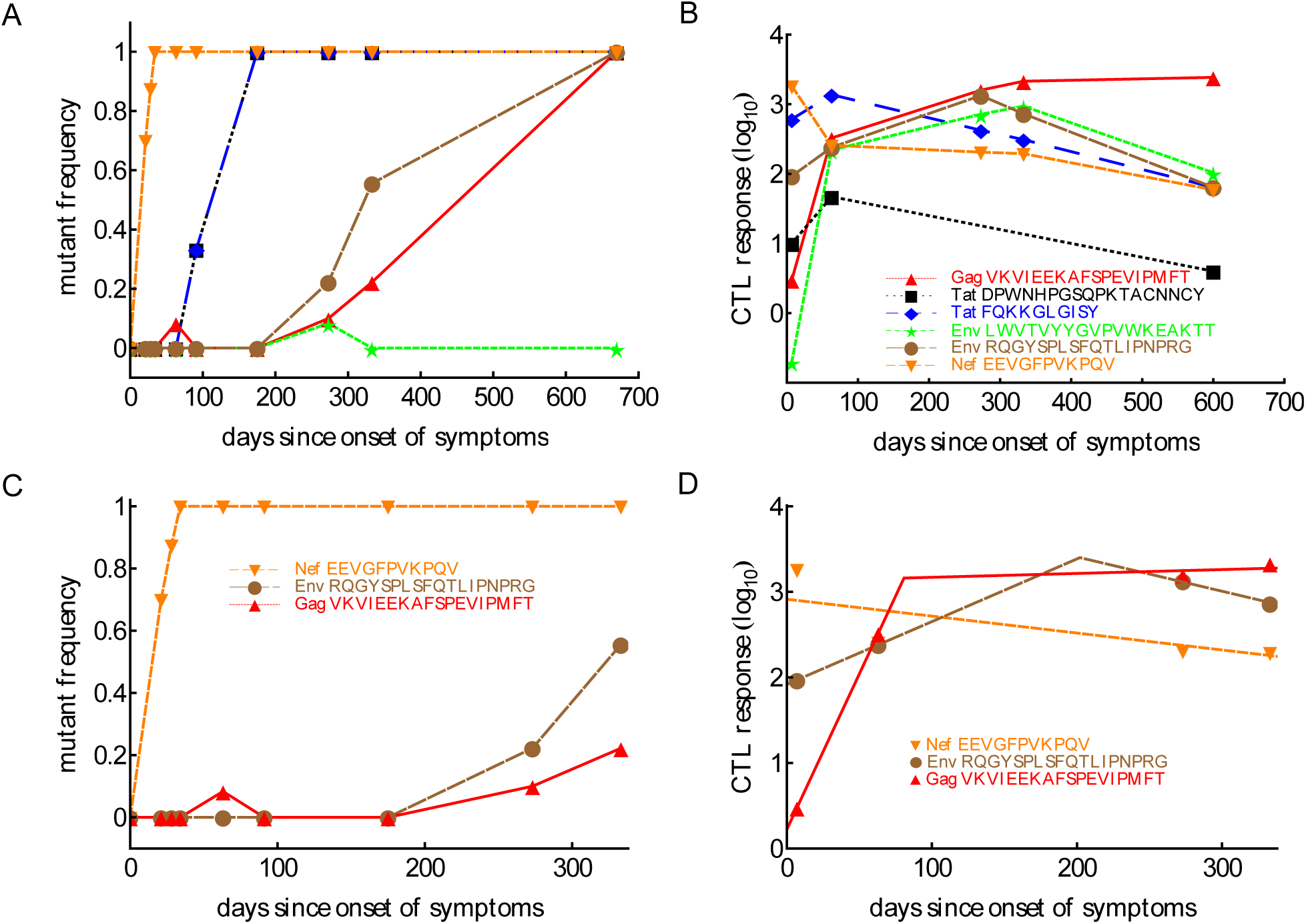
Basic dynamics of CTL response and HIV escape in patient CH131.Patient CH131 had 6 CTL responses (panel B) and 5 responses were escaped by 700 days since infection (panel A). Based on our selection criteria (see Materials and Methods) we focused our analysis on escape in three epitopes: Nef 64-74, Env 709-726 and Gag 156 -173 (panel C) with the corresponding CTL dynamics (panel D). Dashed lines in panel D denote fits of the *T*_on_ - *T*_off_ model (eqn. (6)) to these data resulting in the following estimates for the model parameters for Nef specific T cell responses: *E*_0_ = 808.59 IFN*γ*+ SFC*/*10^6^ PBMC, *α* = 4.55×10^-3^ day^-1^; for Env specific T cell responses: *E*_0_ = 82.97 IFN*γ*+ SFC*/*10^6^ PBMC, *T*_on_ = 0 day, *T*_off_ = 202.02 day, *ρ* = 0.017 day^-1^, *α* = 9.23 × 10^-3^ day^-1^; for Gag specific T cell responses: *E*_0_ = 1.67 IFN*γ*+ SFC*/*10^6^ PBMC, *T*_on_ = 0 day, *T*_off_ = 80.76 day, *ρ*= 0.084 day^-1^, *α* = *-*1.04 × 10^-3^ day^-1^.

We thus fitted 3 different models of viral escape combined with 3 different models for the CTL dynamics to the data on viral escape (Figure 6). Importantly, as with the analysis of data from patient CH159 we found that including the data driven CTL dynamics in the escape models did not improve the quality of the model fit to the escape data (Table 2). In contrast with the previous results, though, the assumption of the constant and time variable killing efficacy (i.e., due to variation in the immune response magnitude) did not strongly impact the quality of the model fit as judged by the AIC or likelihood (Table 2). Importantly, however, models 1&3 gave nearly identical estimates of the CTL killing efficacy, suggesting that for data with good temporal resolution model estimates of the CTL killing efficacy (or by inference, escape rates) are not strongly dependent on the specific mechanisms used to describe escape (independent vs. concurrent escape).

**Table 2:** Parameters estimated by fitting different models of viral escape to escape data in patient CH131 assuming constant killing rates *k*_*i*_ (panels A C), or time varying killing rates due to interpolated CTL response (panels D E) or CTL response in the *T*_on_ - *T*_off_ model (panels G I). Alternative models assume independent escape (model 1, panels A, D, & G), sequential escape (model 2, panels B, E, & H), or concurrent escape (model 3, panels C, F, & I). Fits of models 1&3 gave very close parameter values, but there were some unrealistic parameter values (italicized in the table) from fits of the model 2. *L* and AICc give the log likelihood score and the correlated Akaike information criterion value, respectively. Models 1&3 fit almost equally with three types of response inputs and the lowest *L* and AICc are shown in bold.

|  | peptide | model 1 |  | model 2 |  | model 3 |  |
| --- | --- | --- | --- | --- | --- | --- | --- |
| constant<br>response | | mutation rate<br>($\mu_i, i = 1, 2, 3$) | killing rate<br>($k_i, i = 1, 2, 3$) | mutation rate<br>($\mu_i, i = 1, 2, 3$) | killing rate<br>($k_i, i = 1, 2, 3$) | mutation rate<br>($\mu_i, i = 1, 2, 3$) | killing rate<br>($k_i, i = 1, 2, 3$) |
| | Nef 64-74 | $1.75 \times 10^{-3}$ | 0.25 | $1.72 \times 10^{-3}$ | 0.25 | $1.78 \times 10^{-3}$ | 0.25 |
| | Env 709-726 | $1.03 \times 10^{-7}$ | 0.031 | $3.18 \times 10^{-5}$ | $5.45 \times 10^{-3}$ | $9.91 \times 10^{-7}$ | 0.031 |
| | Gag 156-173 | $1.49 \times 10^{-4}$ | $5.16 \times 10^{-3}$ | *433780.63* | 0.010 | $1.49 \times 10^{-4}$ | $5.19 \times 10^{-3}$ |
| | | $\mathcal{L} = -34.09, AICc = 84.38$ | | $\mathcal{L} = -36.54, AICc = 89.27$ | | $\mathcal{L} = -34.09, AICc = 84.38$ | |
| interpolated<br>response | | mutation rate<br>($\mu_i, i = 1, 2, 3$) | killing rate<br>($k'_i, i = 1, 2, 3$) | mutation rate<br>($\mu_i, i = 1, 2, 3$) | killing rate<br>($k'_i, i = 1, 2, 3$) | mutation rate<br>($\mu_i, i = 1, 2, 3$) | killing rate<br>($k'_i, i = 1, 2, 3$) |
| | Nef 64-74 | $4.33 \times 10^{-4}$ | $1.97 \times 10^{-4}$ | $3.95 \times 10^{-4}$ | $2.00 \times 10^{-4}$ | $4.30 \times 10^{-4}$ | $1.96 \times 10^{-4}$ |
| | Env 709-726 | $7.07 \times 10^{-6}$ | $3.01 \times 10^{-5}$ | $8.76 \times 10^{-5}$ | $1.56 \times 10^{-5}$ | $7.17 \times 10^{-6}$ | $3.00 \times 10^{-5}$ |
| | Gag 156-173 | $1.56 \times 10^{-4}$ | $4.59 \times 10^{-6}$ | *3.33 \times 10^{-3}* | $7.48 \times 10^{-14}$ | $1.55 \times 10^{-4}$ | $4.61 \times 10^{-6}$ |
| | | $\mathcal{L} = -34.02, AICc = 84.24$ | | $\mathcal{L} = -36.79, AICc = 89.77$ | | $\mathcal{L} = -34.02, AICc = 84.24$ | |
| fitted<br>response | | mutation rate<br>($\mu_i, i = 1, 2, 3$) | killing rate<br>($k'_i, i = 1, 2, 3$) | mutation rate<br>($\mu_i, i = 1, 2, 3$) | killing rate<br>($k'_i, i = 1, 2, 3$) | mutation rate<br>($\mu_i, i = 1, 2, 3$) | killing rate<br>($k'_i, i = 1, 2, 3$) |
| | Nef 64-74 | *$3.25 \times 10^{-3}$* | $3.38 \times 10^{-4}$ | *$2.99 \times 10^{-3}$* | $3.46 \times 10^{-4}$ | *$3.16 \times 10^{-3}$* | $3.41 \times 10^{-4}$ |
| | Env 709-726 | $1.38 \times 10^{-6}$ | $2.59 \times 10^{-5}$ | $8.90 \times 10^{-5}$ | $1.05 \times 10^{-5}$ | $1.12 \times 10^{-6}$ | $2.66 \times 10^{-5}$ |
| | Gag 156-173 | $1.73 \times 10^{-4}$ | $2.82 \times 10^{-6}$ | *$3.41 \times 10^{-3}$* | $6.90 \times 10^{-14}$ | $1.73 \times 10^{-4}$ | $2.83 \times 10^{-6}$ |
| | | $\mathcal{L} = -33.94, AICc = 84.07$ | | $\mathcal{L} = -36.98, AICc = 90.17$ | | $\mathcal{L} = -33.94, AICc = 84.07$ | |

**Figure 6:**
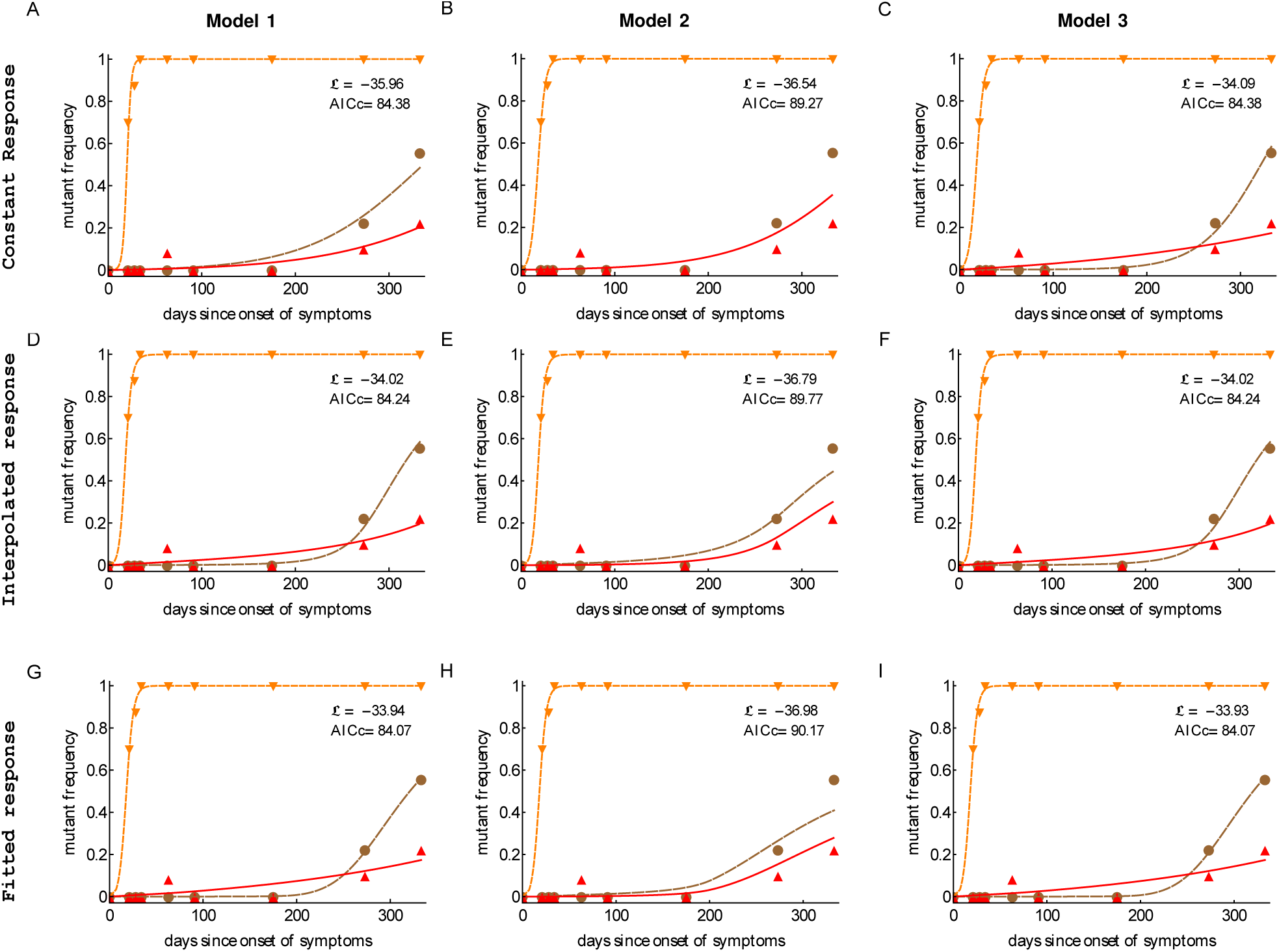
Including CTL response dynamics did not improve model fits of HIV escape data in patient CH131. We fitted model 1 (independent escapes), model 2 (sequential escape) and model 3 (concurrent escape) to escape data in patient CH131 with different CTL response inputs (constant, interpolated or fitted response). Adding data derived time dependent CTL response (interpolated or fitted response) does not improve the fitting results in most cases (Table 2). Notably, model 2 was unable to accurately describe late escape for biologically reasonable mutation rate *µ*_3_. Model parameters providing the best fit are given in Table 2.

Extending the observation made with the patient CH159 data, we found that model assuming sequential escape (model 2) could not accurately describe the dynamics of viral escape for biologically reasonable parameter values specifically for the third escape in Gag156 although this inability was significant only for a constant killing efficacy (Table 2). Allowing time-dependent killing efficacy resulted in small yet larger values for the mutation rate than that expected from basic calculations. Forcing the mutation rate *µ*_3_ to be constrained (*µ*_3_ ≤ 10^-3^) significantly reduced the quality of themodel fit to data (likelihood ratio test, *p*≪0.001). Furthermore, estimates for the CTL killing efficacy differed between model 2 and models 1&3 suggesting that model choice (sequential vs. concurrent) may indeed influence estimates of the killing efficacy.

### 3.3 No difference in predicted killing efficacy of CTLs, specific to different epitopes

Our analyses so far demonstrated that several different mathematical models were capable of accurately describing the escape data, but this ability was dependent on the specific pathway of how escape mutants were generated and the assumption on whether data-driven CTL dynamics was included in the model. In cases, when a model was able to accurately describe the data, we generally observed different estimates for the parameters for HIV escape in different epitopes; for example, for the data in patient CH131 estimated CTL killing rate in the model 1 (independent escapes) with interpolated response different nearly 100 fold between 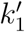 and 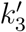 (Table 2). Knowing which immuneresponses may be more efficient on a per cell basis in killing virus-infected cells may be beneficialfor inducing such responses by vaccination. We therefore investigated how robust these differences in estimated per capita killing rates are. For that we fitted mathematical models assuming equal killing efficacies to the data on escape. As expected, reducing the number of fitted parameters led to fits of lower quality (as judged by the log likelihood); however, this reduction in complexity of the model was favored by the AIC and in most cases by the likelihood ratio test (Tables S2 and S4 in Supplement). Visually, the reduction in the quality of the model fit to data was also relatively small (Figures S2 and S4 in Supplement). Thus, for these data we found no strong evidence in the difference in the estimated per capita killing efficacy of the CTL response specific to different viral epitopes.

### 3.4 Identifying conditions when the model 2 (sequential escapes) fails

In analysis of data from both patients we found that model 2, describing sequential escape from CTL responses, was not able to accurately describe experimental data for biologically reasonable parameter values; these model fits predicted extremely high mutation rates (e.g., see Tables 1 and 2). Additional analyses demonstrated that fitting the models with constrained mutation rates, µ_*i*_ ≤10^-3^,led to fits of significantly lower quality (based on increased AIC, results not shown).

A closer look at the experimental data for which model 2 provided unreasonably high mutation rates revealed that the trajectories of two subsequent escapes in the model 2 were too close to each other which naturally required a high mutation rate from one variant to another. Therefore, only when trajectories are separated in time mutation rate *µ*_2_ is expected to be biologically reasonable. Indeed, by simulating virus dynamics using model for sequential escapes by varying model parameters we found that CTL killing rate has the major impact on the time delay between two escapes (Figure 7). This analysis thus suggested that for the model 2 (sequential escape) to be consistent with the data, escapes from 2 responses must be separated in time by about 20 50 days.

**Figure 7:**
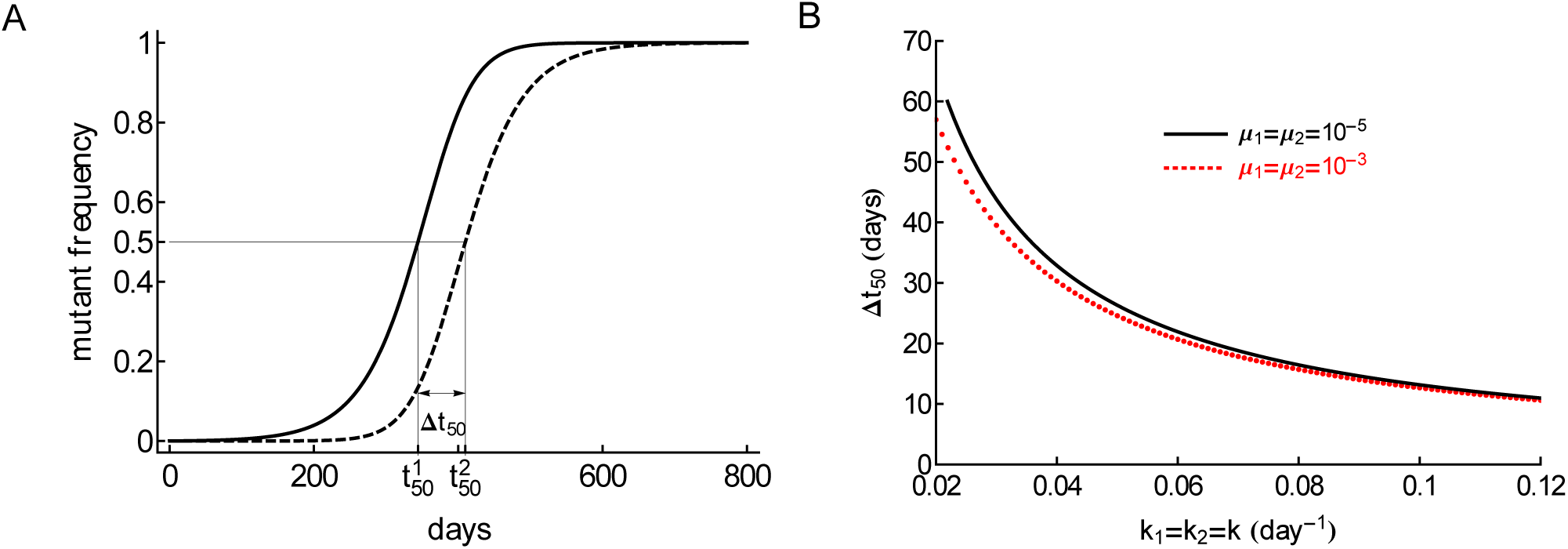
Model, assuming sequential escape (model 2), can be consistent with escape data when the trajectories for two sequential viral escape are separated in time. We illustrate that separation of trajectories by Δ*t*_50_ = 409.8 – 344.2 ≃ 66 days is sufficient for the mutation rate to be realistically small (panel A). Here*t*^*i*^_50_ is the time by which the *i*^*th*^ variant reaches 50% of the viral population, so Δ*t*_50_ = *t*^2^_50_ – *t*^1^_50_. Parameters used in simulations are *µ*_1_ = *µ*_2_ = 10^-5^, *k*_1_ = *k*_2_ = 0.02 day^-1^, *r* = 1.5 day^-1^, *δ* = 1 day^-1^. The distance between trajectories needed for small predicted mutation rates is reduced for higher CTL killing rates (panel B) and the time is only weakly dependent on the mutation rate assumed in simulations.

## 4 Discussion

CTL responses play a major role in HIV within-host evolution [36, 37]. Recent studies suggested that a relative magnitude of the CTL response (relative immunodominance) plays an important role in determining the time of viral escape from T cell responses [4, 31]. These previous studies, however, only utilized a maximum value of the CTL response early in infection, in general within 50 days since the onset of symptoms, and thus impact of the kinetics of CTL response on the rate of virus escape remained undetermined. Furthermore, the pathways of HIV escape from CTL responses were not fully resolved as escapes occurring sequentially and concurrently have been proposed [26, 29, 39], and several previous studies assumed that escapes occur independently from each other [2, 12, 16]. Here by using experimental data on evolution of HIV sequences from acute infection into chronic phase and temporally resolved dynamics of HIV-specific CTL responses we tested the hypothesis that CTL dynamics plays an important role in virus escape.

Perhaps in contrast with our initial expectations (e.g., due to [4, 31]), we found that including experimentally measured dynamics of epitope specific CTL responses did not led to a better description of the kinetics of viral escape from T cells (e.g., in patient CH131, Table 2), or even reduced the quality of the model for viral escape fit to data (e.g., in patient CH159, Table 1). This was not because we assumed that killing of virus infected cells was dependent on the absolute magnitude of epitope specific CTL responses; assuming frequency dependent killing, that is, when killing of infected cells expressing *i*^*th*^ epitope was given by 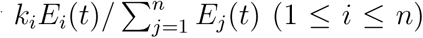 led to similar conclusions (results not shown). Because previous work suggested that kinetics of escape was in dependent of the specific mechanism of how CTLs suppress wild-type virus (e.g., killing of infected cells or virus production by infected cells) [15], we did not investigate non lytic control of HIV by T cells. It is interesting that the lack of correlation between the rate of viral escape and CTL response magnitude was highlighted previously [16].

Reasons of why a model with time variable CTL response did not describe experimental data better than a model with a constant response remain unclear but several hypotheses could be generated. First, frequency of sampling of the viral sequences may not be high enough to detect change in the speed at which mutant viruses accumulate in the population. Indeed, in mathematical models CTL dynamics has a direct impact on the rate of escape (e.g., see eqn. (3)) and the observed changes in CTL densities may not be reflected in escape data if data sampling is infrequent. Second, virus sequence data could simply be noisy. Because only handful of viral sequences were analyzed by the SGA/S, measurements of frequencies of viral variants have in general large expected error (e.g., Figure 2). Third, CTL dynamics in the blood may not reflect CTL dynamics in tissues such as secondary lymphoid organs (lymph nodes and spleen). While it is well known that T cells recirculate in the body [13], how quickly CTLs in the tissues migrate into the blood and then back to the tissues during HIV infection is not known. Finally, it is possible that the measured CTL responses were not the drivers of escape. While the ability of CTLs to recognize the wild-type virus and inability of the same CTLs to recognize mutant viruses is generally interpreted as evidence that these CTLs drove viral escape, such observations are correlational in nature, and thus can not fully establish the causality of escape, at least in humans.

Our results may be interpreted as contradictory to several previous studies that found a strong correlation between the time of viral escape (time when a escape variant reaches frequency of 50% in the viral population) and a relative magnitude of CTL response (relative or “vertical” immunodom inance) [4, 31]. However, our studies are not directly compatible because this previous work focused on the timing of escape while we primarily focused on the rate of viral escape. These two parameters are differently impacted by the CTL response [16] and may have different clinical importance. In our simple mathematical model (e.g., eqn. (3)) CTL response magnitude is expected to directly impact the rate at which an escape mutant accumulates in the population, independently of when this escape may occur. In contrast, timing of viral escape also depends on the mutation rate. Biologically, however, timing of escape may be more important than the rate because it may be more beneficial to the patient if viral escape occurs 5 years after infection but rapidly as compared to slow escape in just 1 year. This conjecture clearly depends on the premise that HIV escapes from CTL responses are detrimental to patients.

In our analysis we generally found that for well sampled data the pathway of generation of escape mutants played a minor role in predicting overall CTL killing efficacy; assuming escapes that occur independently (model 1) or concurrently (model 3) gave nearly identical estimates of the CTL killing efficacy (e.g., Tables 1 and 2). In contrast, the model assuming sequential escape (model 2) often failed to accurately explain experimental data; this was due to some escapes co-occurring at nearly the same time which obviously violated the model assumption of sequential escape. This inability of the sequential escape model to describe the data may be the result of the way we compared models to data: by using deterministic model approach and by ignoring recombination. Using deterministic model may be justified because in acute infection the effective population size of HIV may be sufficiently large and ignoring recombination may again be appropriate because very few cells in HIV infection are generally infected by 2 or more viruses [24, 25]. However, further work is needed to demonstrate whether our conclusions regarding inability of sequential escape model to accurately explain some escape data is due to some of the assumptions made in the model by running stochastic simulations and by allowing some degree of recombination.

Many of our model fits predicted a high mutation rate for the first epitope to be escaped by thevirus (e.g., Table 2). This model prediction could not be changed by shifting the experimental data to allow for more time to generate escape mutant; in part, this test failed because in the absence of epitope specific T cells escape variants accumulate rather slowly mainly driven by mutations. It may indicate that immune pressure on the virus population starts much earlier than it is reflected in the blood, echoing our concerns of whether CTL dynamics in the blood is an accurate reflection of T cell response in lymphoid tissues. Currently it is believed that lymphoid tissues and not the blood are the major places of interactions between the virus and CTLs [22, 30].

Our analysis further highlights the importance of choosing the appropriate statistical model for the analysis of the escape data – assuming normally-distributed residuals, and therefore, using least squares approach, may not be appropriate for some escape data with very few sequences analyzed. Importantly, we confirm that the type of statistical model has an impact on the estimate of the escape rate [17].

We found that experimental data on HIV escape can be explained well if we assume identical per capital killing efficacy of CTLs, specific to different viral epitopes. This suggests that individual per capita killing rates not accurately estimated from these data. While it is possible that this result was the consequence of assuming additive killing of virus infected cells by different CTL responses, we currently do not have any *in vivo* data to support more complex killing terms.

Overall, analyses of data from two patients suggested that models assuming independent escape of HIV from different CTL responses (model 1) or models assuming concurrent escape from mul tiple CTL responses (model 3) fit the data well and provide very similar (often nearly identical) estimates for the killing efficacy of CTL response. Thus, for well sampled data assumption of in dependent escapes may be sufficient to accurately estimate HIV escape rates. Also the model with data driven time dependent CTL response (interpolated or fitted response input) did not improve the quality of the model fit to data, so at present it appears to be unnecessary to incorporate the experimentally-measured CTL response dynamics in the model describing viral escapes. Our analysis thus demonstrates how mathematical modeling may help to quantify HIV evolution in presence of CTL responses and to highlight potential limitations with experimental measurements.

## Author Contributions

YY and VG designed the study. YY performed the simulations. YY and VG contributed to analysis, interpolation of data and simulation results. YY and VG wrote the paper.

## Acknowledgments

The authors would like to thank Dr. Nilu Goonetilleke and the Center for HIV/AIDS Vaccine Immunology (CHAVI) for data access.sss

## Supplementary Information

### S1 Model derivation of viral escape from multiple CTL responses

Following the previous work [16], we use *m***_i_** to denote the density of variants denoted by a vector **i** = (*i*_1_, *i*_2_, *…, i*_*n*_), which is the index denoting the positions of *n* epitopes, and we define *i*_*j*_ = 0 if there is no mutation in the *j*^*th*^ CTL epitope and *i*_*j*_ = 1 if there is a mutation leading to an escape from the *j*^*th*^ CTL response.

We assume that a CTL response that recognizes the *i*^*th*^ epitope of the virus kills the virus infected cells at rate *k*_*i*_, and escaping from the *i*^*th*^ CTL responses only at a rate *µ*_*i*_ leads to a viral replicative fitness cost *c*_*i*_ (*i* = 1, *…, n*). As shown in model (1) of viral escape from a single CTL response (see equation (1)), we denote the infection rate of variants *m***_i_** by *β***_i_** and variants *m***_i_** are produced by infected cells at rate *p***_i_** (**i** ∈ *I*). We assume that the wild-type has a higher (or equal) reproductive ratio, that is *β***_i_***p***_i_** ≤ *β*_(0,0,*…,*0)_*p*_(0,0,*…,*0)_ for all **i** ≠ (0, 0,…, 0) (i ∈ *I*).

Let 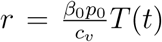 (with *β*_0_ = *β*_(0,0,*…,*0)_ and *p*_0_ = *p*_(0,0,*…,*0)_) as the reproduction rate of wild-type virus, we use fitness cost *c*_*i*_ (*i* = 1, *…, n*) and *r* to express the replication rate of each escape variant. For simplicity, we neglect recombination and only allow single point mutation. To be consistent with the model of viral escape from a single CTL escape, we let *β*_*i*_ denote the rate at which variant *m*_(0,*…,*1,*…,*0)_ (only *i*^*th*^ position equal to 1) infect cells, and *p*_*i*_ denote the production rate of variant *m*_(0,*…,*1,*…,*0)_. Then the fitness cost *c*_*i*_ of *m*_(0,*…,*1,*…,*0)_ can be written as 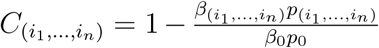. As for variants *m*_(*i*_1,*…,in*) having two more mutations, we assume

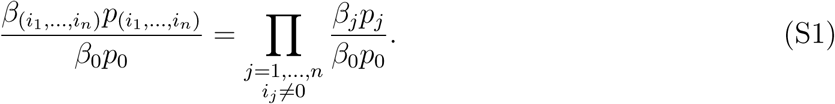

This assumption means for variant having mutations at *i*^*th*^ and *j*^*th*^ epitopes, the normalized reproduc tive rate (by wild-type reproductive rate *β*_0_*p*_0_) equals the product of normalized reproductive rates of variants, which only have one mutation at *i*^*th*^ or *j*^*th*^ epitope. For example, 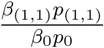=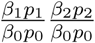 with *n* = 2. Under this assumption, the fitness cost 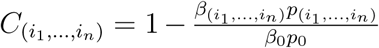 of variant *m*(*i*_1_,*…,i*_*n*_) can be written as

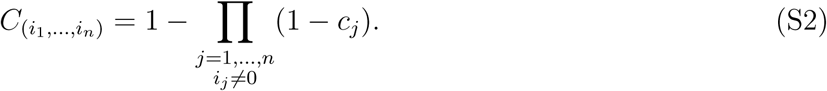

Assuming multiplicative fitness, the fitness cost of a variant **i** = (*i*_1_, *i*_2_, *…, i*_*n*_) is 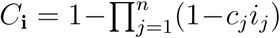. The death rate of the escape variant **i** = (*i*_1_, *i*_2_, *…, i*_*n*_) due to remaining CTL responses is given by 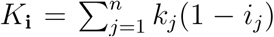, where we assume that killing of infected cells by different CTL responses is additive.

We neglect recombination and backward mutation from mutant to wild-type in this modeling framework. More specifically, for two escape variants *m***_i_** = *m*_(*i*_1_,*i*_2_,…,*i_n_*)_ and *m***_j_** = *m*_(*j*_1_,*j*_2_,…,*j_n_*)_, we define the mutation rate *M***_*i,j*_** from *m***_i_** to *m***_j_** as *µ*_*k*_, if and only if *m***_j_** has only one more mutation at position *k* than *m***_i_** and all other positions are exactly same. For example, when there are 3 CTL responses, the mutation rate from *m*_(1,0,0)_ to *m*_(1,0,1)_ is *µ*_3_, and the mutation rate from *m*_(0,0,0)_ to *m*_(1,0,1)_ is 0.

Similar as equation (1), the dynamics of the wild-type and all escapes from CTL responses is given by

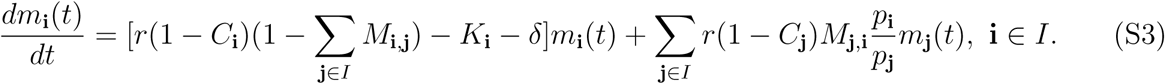

Here we adopt the simple assumption that escape mutants and wild-type viruses may differ from rates *β*_**i**∈*I*_ at which they infect cells, that is *p*_0_ = *p***_i_** and *β*_0_ ≥ *β***_i_** (**i** ∈ *I* and **i** ≠ (0, *…*, 0)). The system (S3) becomes

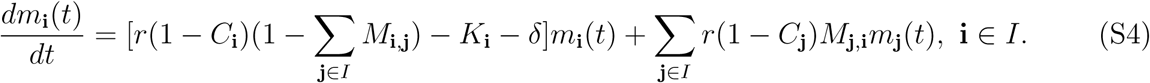

We define 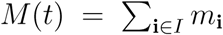 as the total density of all variants in the population, and *f*_*j*_(*t*) (*j* = 1, *…, n*) is the fraction of viral variants that have escaped recognition from the *j*^*th*^ CTL response. Then

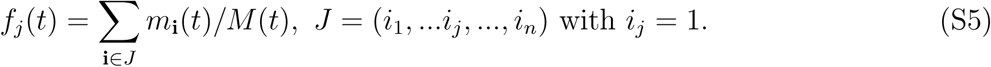

For example, when *n* = 2, there are 3 types of escape variants *m*_(0,0)_, *m*_(1,0)_ and *m*_(1,1)_ for “sequential” escape (model 2), and 4 types of escape variants *m*_(0,0)_, *m*_(1,0)_, *m*_(0,1)_ and *m*_(1,1)_ for “concurrent” escape (model 3).

Under all above assumptions, from system (S4), model 2 with *n* = 2 can be written as:

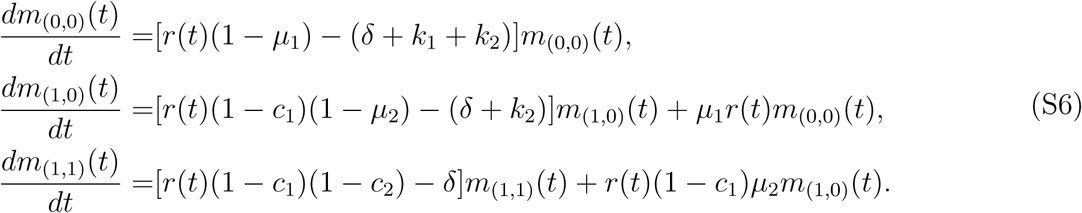

and

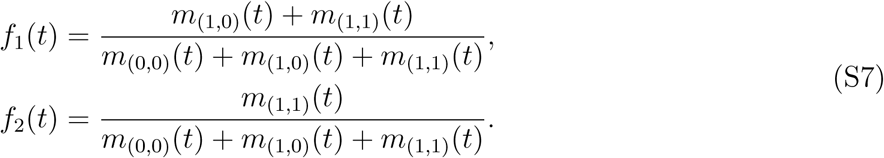

Similarly, following system (S4), model 3 with *n* = 2 can be written as:

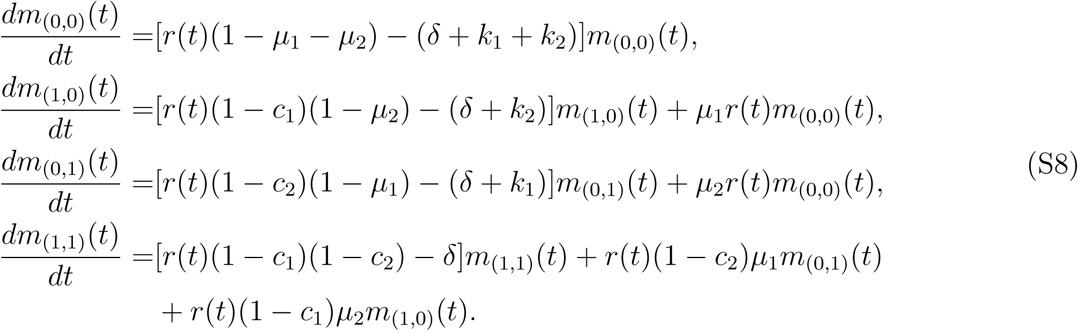

and

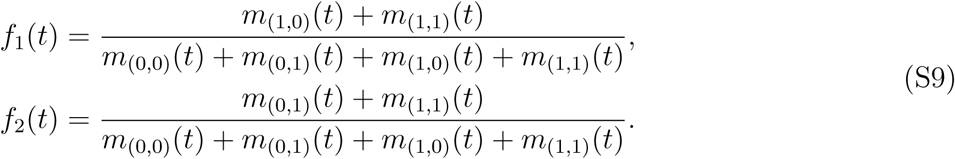

### S2 Examples of “sequential” and “concurrent” escapes for *n* = 3 epi topes/CTL responses

The difference between “sequential” escape (model 2) and “concurrent” escape (model 3) is the set of escape variants *I*. The set *I* has *n* + 1 elements for “sequential” escape model and 2^*n*^ elements for “concurrent” escape model for *n* epitope case. For the simple case *n* = 3, equations for all escape variants are

**Model 2:**

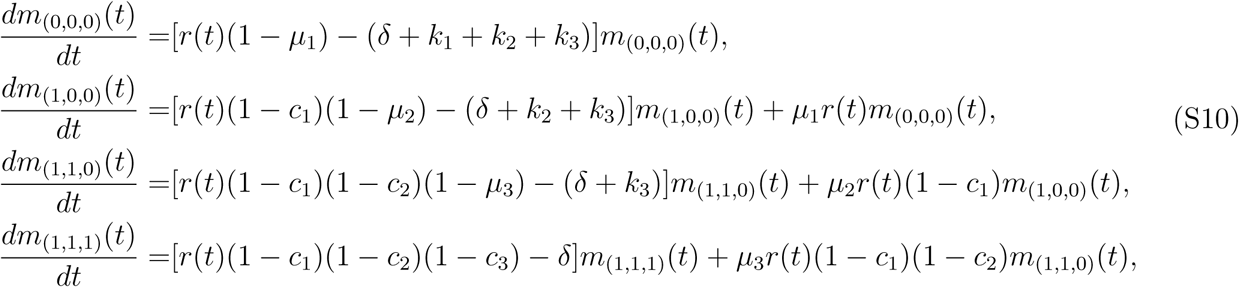

and

**Model 3:**

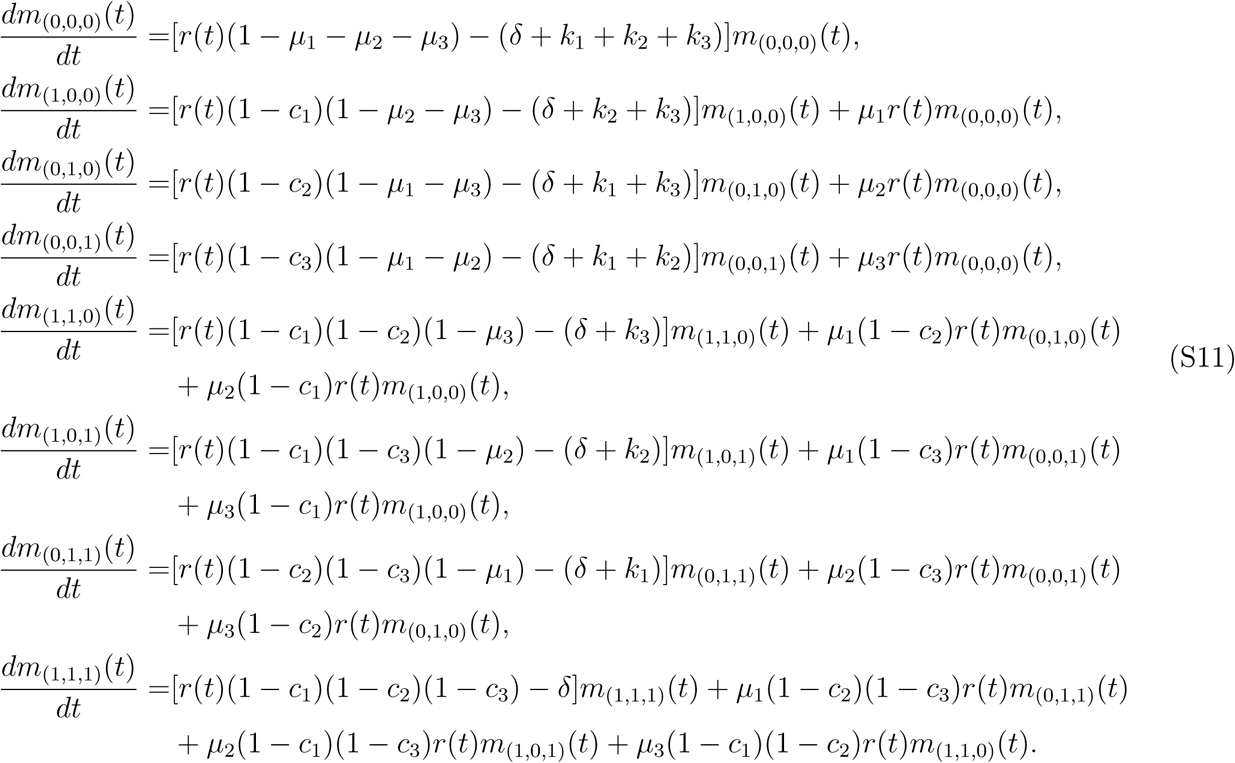

### S3 Additional results of the analysis

**Figure S1:**
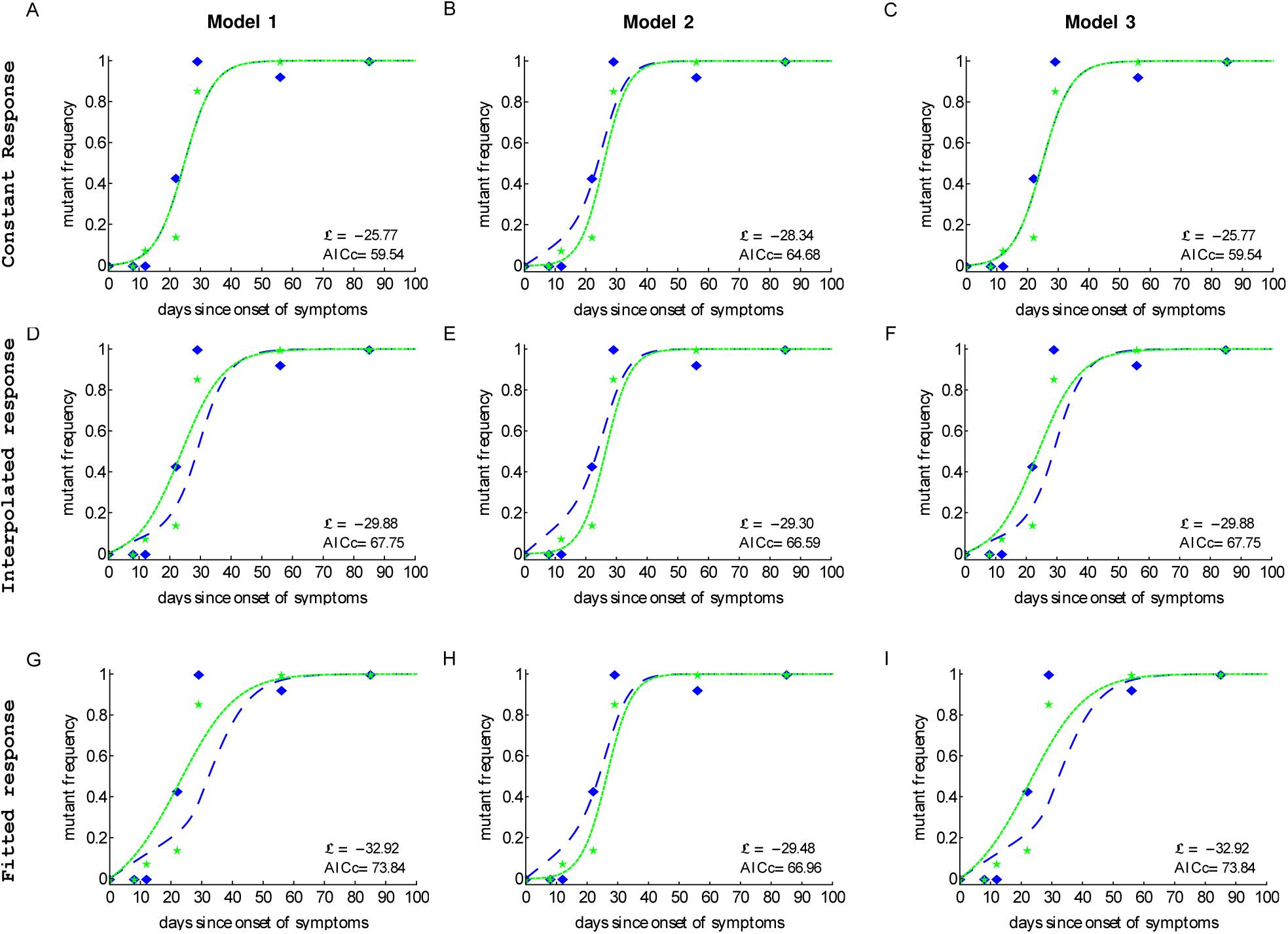
Mathematical model accurately explains kinetics of HIV escape from CTL response when assuming equal mutation rates (*µ*_1_ = *µ*_2_) for data from patient CH159. We fit the three mathematical models (models 1, 2, and 3) to experimental data using likelihood approach outlined in the Materials and Methods section assuming *µ*_1_ = *µ*_2_. Three different models for the CTL response dynamics were assumed: constant input, interpolated input and fitted input. Models with response input did not improve the quality of the model fit to data. The best fit was provided by the models 1&3 with constant response. Estimated parameter values are given in Table S1. Notations for data points and lines are identical to those given in Figure 4 in the main text.

**Table S1:** Best fit parameters of three models (models 1, 2 and 3) fitted to experimental data on HIV escape in patient CH159 assuming identical mutation rates (*µ*_1_ = *µ*_2_). Model fits are shown in Figure S1. ℒ and AICc give the log likelihood score and the correlated Akaike information criterion value, respectively. Best (maximum) and AICc ℒ (minimum) scores are shown in bold. Mutation rates which exceed a theoretically assumed maximum value of 10^-3^ are shown in italics.

|  | peptide | model 1 |  | model 2 |  | model 3 |  |
| --- | --- | --- | --- | --- | --- | --- | --- |
| | | mutation rate<br>($\mu_i$, i=1,2) | killing rate<br>($k_i$, i=1,2) | mutation rate<br>($\mu_i$, i=1,2) | killing rate<br>($k_i$, i=1,2) | mutation rate<br>($\mu_i$, i=1,2) | killing rate<br>($k_i$, i=1,2) |
| constant<br>response | Rev 65-82 | $7.55 \times 10^{-4}$ | 0.21 | $6.75 \times 10^{-3}$ | $1.86 \times 10^{-13}$ | $7.55 \times 10^{-4}$ | 0.21 |
|  | Nef 177-194 |  | 0.21 |  | 0.25 |  | 0.21 |
| | | $\mathcal{L} = -\mathbf{25.77}$, $AICc = \mathbf{59.54}$ | | $\mathcal{L} = -28.34$, $AICc = 64.68$ | | $\mathcal{L} = -\mathbf{25.77}$, $AICc = \mathbf{59.54}$ | |
| interpolated<br>response | | mutation rate<br>($\mu_i$, i=1,2) | killing rate<br>($k'_i$, i=1,2) | mutation rate<br>($\mu_i$, i=1,2) | killing rate<br>($k'_i$, i=1,2) | mutation rate<br>($\mu_i$, i=1,2) | killing rate<br>($k'_i$, i=1,2) |
| | Rev 65-82 | $5.98 \times 10^{-3}$ | $3.07 \times 10^{-3}$ | $8.88 \times 10^{-3}$ | $3.69 \times 10^{-12}$ | $4.98 \times 10^{-3}$ | $3.07 \times 10^{-3}$ |
| | Nef 177-194 | | $1.51 \times 10^{-3}$ | | $2.74 \times 10^{-3}$ | | $1.51 \times 10^{-3}$ |
| | | $\mathcal{L} = -29.88$, $AICc = 67.75$ | | $\mathcal{L} = -29.30$, $AICc = 66.59$ | | $\mathcal{L} = -29.88$, $AICc = 67.75$ | |
| fitted<br>response | | mutation rate<br>($\mu_i$, i=1,2) | killing rate<br>($k'_i$, i=1,2) | mutation rate<br>($\mu_i$, i=1,2) | killing rate<br>($k'_i$, i=1,2) | mutation rate<br>($\mu_i$, i=1,2) | killing rate<br>($k'_i$, i=1,2) |
| | Rev 65-82 | $8.73 \times 10^{-3}$ | $1.98 \times 10^{-3}$ | $8.19 \times 10^{-3}$ | $6.32 \times 10^{-9}$ | $8.72 \times 10^{-3}$ | $1.98 \times 10^{-3}$ |
| | Nef 177-194 | | $9.22 \times 10^{-4}$ | | $2.69 \times 10^{-3}$ | | $9.23 \times 10^{-4}$ |
| | | $\mathcal{L} = -34.92$, $AICc = 77.85$ | | $\mathcal{L} = -29.48$, $AICc = 66.97$ | | $\mathcal{L} = -34.92$, $AICc = 77.85$ | |

**Figure S2:**
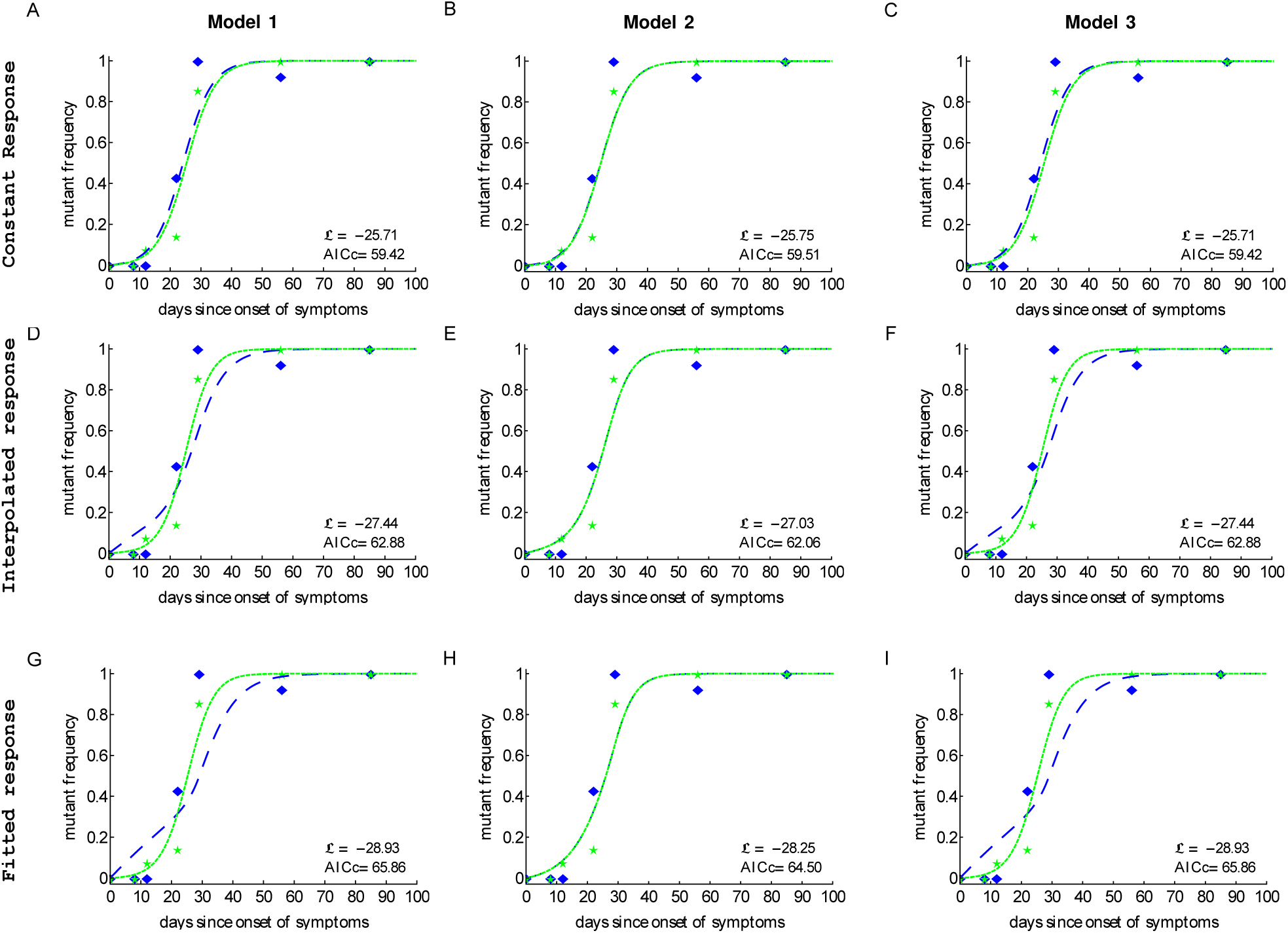
Killing rates of CTL responses specific to different epitopes may be similar. We fit three different models to experimental data from patient CH159 assuming identical CTL killing rates (*k*_1_ = *k*_2_) with different CTL response dynamics (constant input, interpolated input and fitted input). Such constrain did not reduce the quality of the model fit to data as judged by AIC. Parameter estimates are given in Table S2. Notations for data points and lines are identical to those given in Figure 4 in the main text.

**Table S2:**
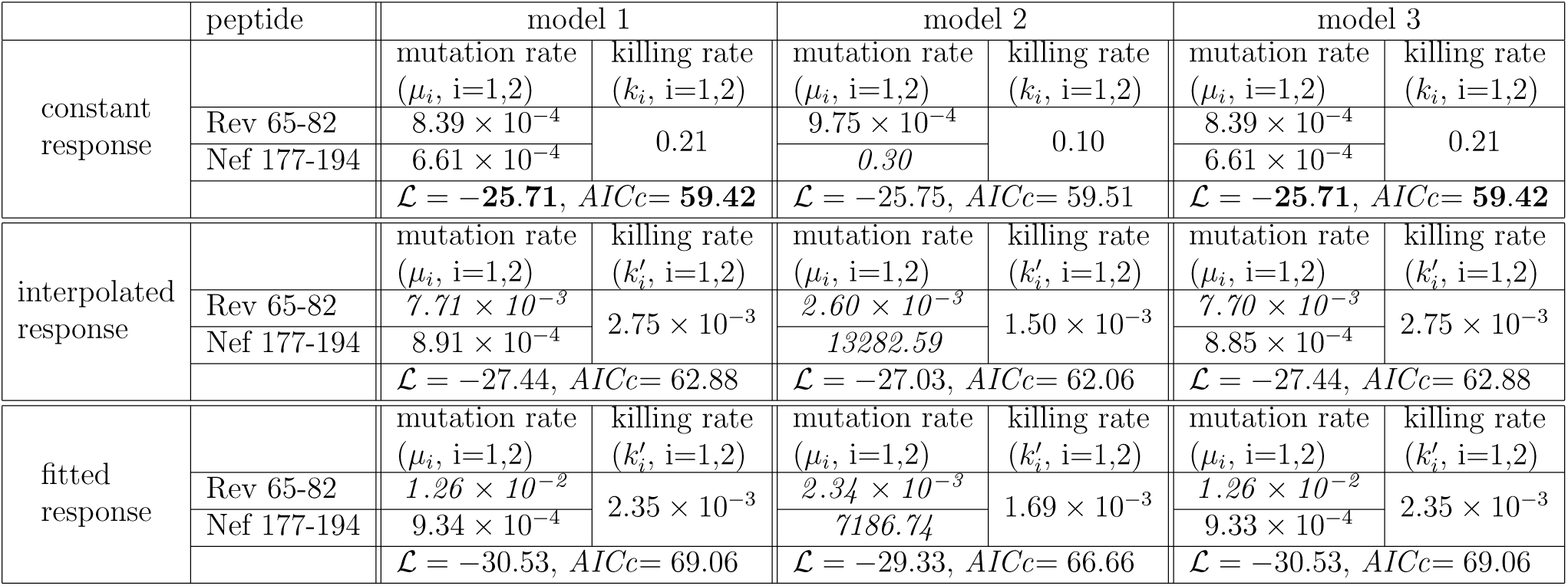
Best fit parameters of three models (models 1, 2 and 3) fitted to experimental data on HIV escape in patient CH159 assuming identical killing rates (*k*_1_ = *k*_2_). Model fits are shown in Figure S2. High (perhaps unrealistic) mutation rates are highlighted in italic. ℒ and AICc give the log likelihood score and the correlated Akaike information criterion value, respectively. Best ℒ (maximum) and AICc (minimum) scores are shown in bold.

**Figure S3:**
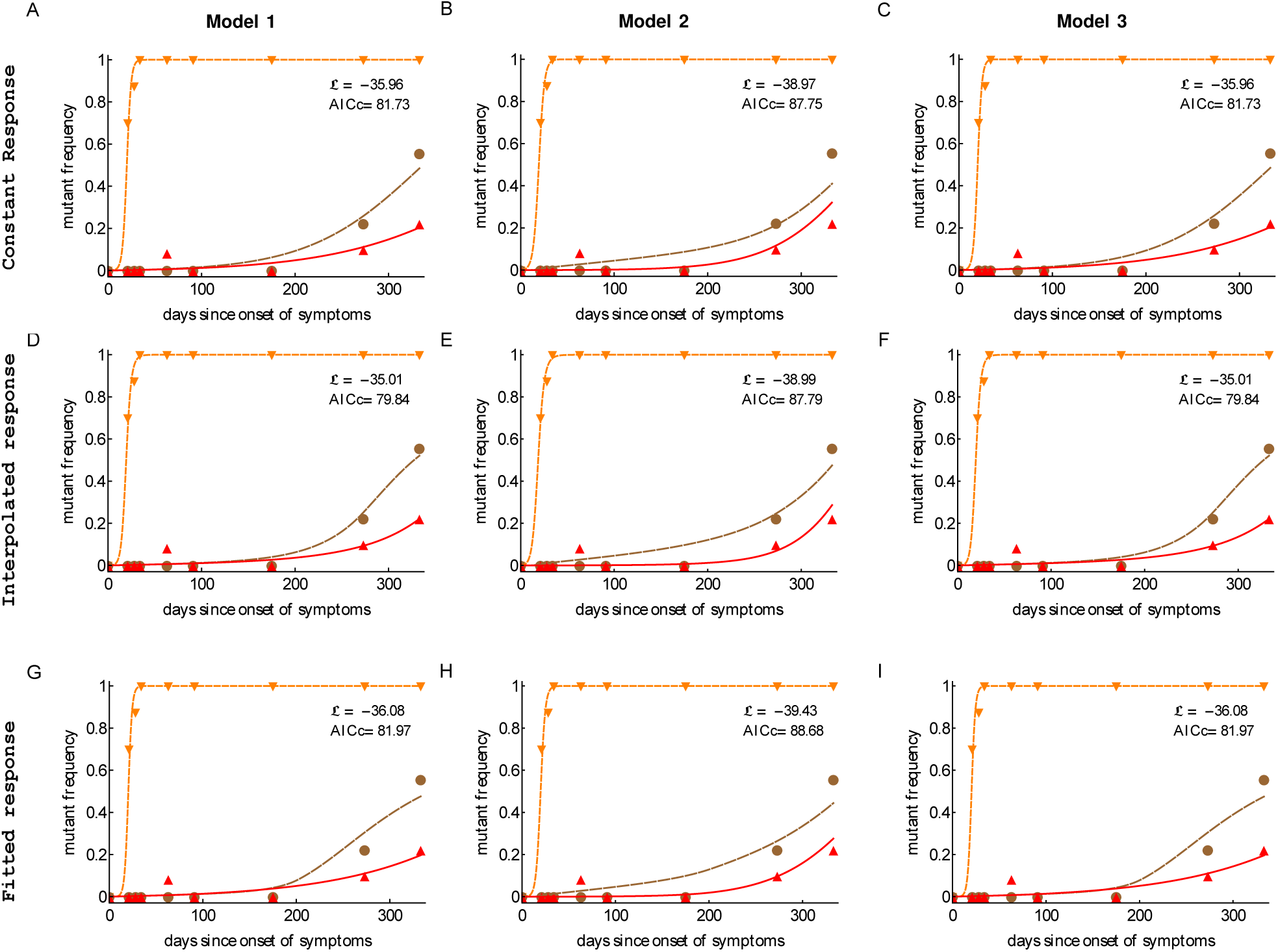
Mathematical models accurately explain kinetics of HIV escape from CTL response when assuming equal mutation rates (*µ*_1_ = *µ*_2_ = *µ*_3_) for data from patient CH131. We fit the three mathematical models (models 1, 2, and 3) to experimental data using likelihood approach outlined in the Materials and Methods section assuming *µ*_1_ = *µ*_2_ = *µ*_3_. Three different models for the CTL response dynamics were assumed: no input, interpolated input and fitted input. Best model fit was provided by the models 1&3 with interpolated response input. Estimated parameter values are given in Table S3. Notations for data points and lines are identical to those given in Figure 6 in the main text.

**Table S3:** Best fit parameter values found by fitting different mathematical models (models 1, 2 and 3) to experimental data in patient CH131 assuming identical mutation rates (*µ*_1_ = *µ*_2_ = *µ*_3_). Model fits are shown in Figure S3. ℒ and AICc give the log likelihood score and the correlated Akaike information criterion value, respectively. Best ℒ (maximum) and AICc (minimum) scores are bolded in the table.

|  | peptide | model 1 |  | model 2 |  | model 3 |  |
| --- | --- | --- | --- | --- | --- | --- | --- |
| Constant<br>response | | mutation rate<br>($\mu_i, i = 1, 2, 3$) | killing rate<br>($k_i, i = 1, 2, 3$) | mutation rate<br>($\mu_i, i = 1, 2, 3$) | killing rate<br>($k_i, i = 1, 2, 3$) | mutation rate<br>($\mu_i, i = 1, 2, 3$) | killing rate<br>($k_i, i = 1, 2, 3$) |
| | Nef 64-74 | $4.34 \times 10^{-5}$ | 0.44 | $3.05 \times 10^{-4}$ | 0.34 | $4.34 \times 10^{-5}$ | 0.44 |
| | Env 709-726 | | 0.016 | | $1.44 \times 10^{-10}$ | | 0.016 |
|  | Gag 156-173 |  | 0.011 |  | 0.021 |  | 0.011 |
| | | $\mathcal{L} = -35.96, AICc = 81.73$ | | $\mathcal{L} = -38.97, AICc = 87.75$ | | $\mathcal{L} = -35.96, AICc = 81.73$ | |
| interpolated<br>response | | mutation rate<br>($\mu_i, i = 1, 2, 3$) | killing rate<br>($k'_i, i = 1, 2, 3$) | mutation rate<br>($\mu_i, i = 1, 2, 3$) | killing rate<br>($k'_i, i = 1, 2, 3$) | mutation rate<br>($\mu_i, i = 1, 2, 3$) | killing rate<br>($k'_i, i = 1, 2, 3$) |
| | Nef 64-74 | $6.22 \times 10^{-5}$ | $2.53 \times 10^{-4}$ | $3.00 \times 10^{-4}$ | $2.07 \times 10^{-4}$ | $6.22 \times 10^{-5}$ | $2.53 \times 10^{-4}$ |
| | Env 709-726 | | $1.87 \times 10^{-5}$ | | $5.33 \times 10^{-6}$ | | $1.87 \times 10^{-5}$ |
| | Gag 156-173 | | $8.73 \times 10^{-6}$ | | $1.24 \times 10^{-5}$ | | $8.73 \times 10^{-6}$ |
| | | $\mathcal{L} = -\mathbf{35.01}, AICc = \mathbf{79.84}$ | | $\mathcal{L} = -38.99, AICc = 87.79$ | | $\mathcal{L} = -\mathbf{35.01}, AICc = \mathbf{79.84}$ | |
| fitted<br>response | | mutation rate<br>($\mu_i, i = 1, 2, 3$) | killing rate<br>($k'_i, i = 1, 2, 3$) | mutation rate<br>($\mu_i, i = 1, 2, 3$) | killing rate<br>($k'_i, i = 1, 2, 3$) | mutation rate<br>($\mu_i, i = 1, 2, 3$) | killing rate<br>($k'_i, i = 1, 2, 3$) |
| | Nef 64-74 | $7.48 \times 10^{-5}$ | $6.72 \times 10^{-4}$ | $3.21 \times 10^{-4}$ | $5.41 \times 10^{-4}$ | $7.65 \times 10^{-5}$ | $6.70 \times 10^{-4}$ |
| | Env 709-726 | | $1.19 \times 10^{-5}$ | | $2.88 \times 10^{-6}$ | | $1.18 \times 10^{-5}$ |
| | Gag 156-173 | | $5.82 \times 10^{-6}$ | | $1.00 \times 10^{-5}$ | | $5.74 \times 10^{-6}$ |
| | | $\mathcal{L} = -36.08, AICc = 81.97$ | | $\mathcal{L} = -39.43, AICc = 88.68$ | | $\mathcal{L} = -36.08, AICc = 81.97$ | |

**Figure S4:**
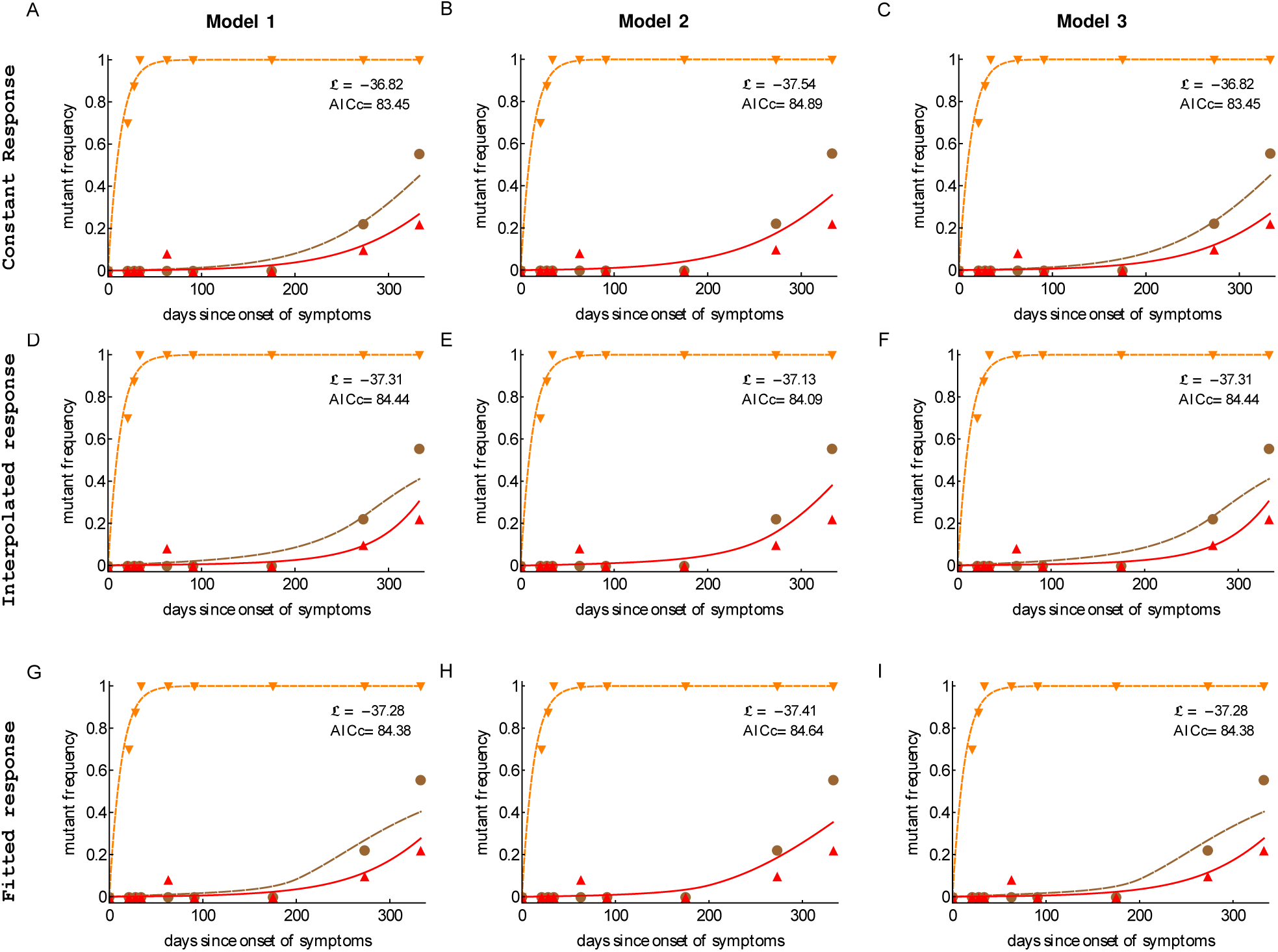
No evidence for difference in CTL killing rates for data on HIV escape in patient CH131. We fit different mathematical models (models 1, 2, or 3) to experimental data from patient CH131 assuming equal killing rates (*k*_1_ = *k*_2_ = *k*_3_). The best fit is given by models 1&3 with interpolated response input. Best fit parameter values are given in Table S4. Notations for data points and lines are identical to those given in Figure 6 in the main text.

**Table S4:**
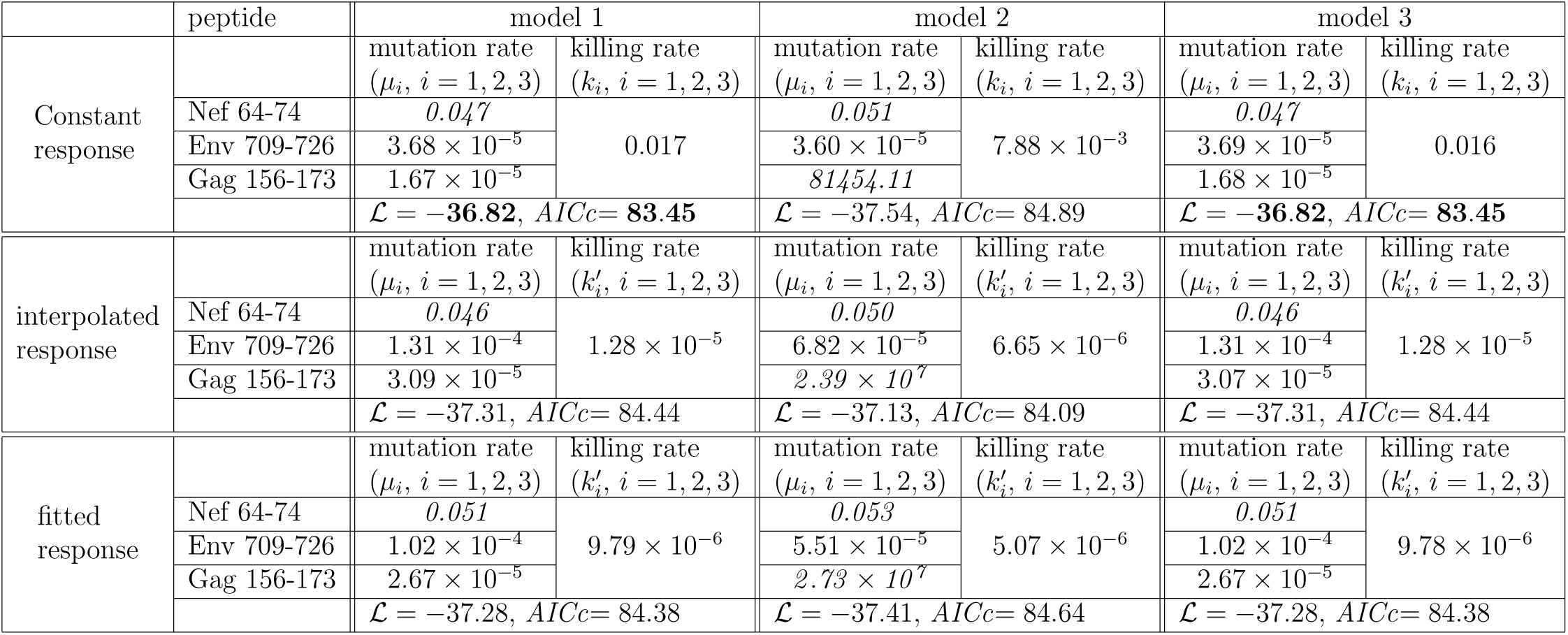
Best fit parameter values found by fitting different mathematical models (models 1, 2 and 3) to experimental data in patient CH131 assuming identical killing rates (*k*_1_ = *k*_2_ = *k*_3_). Model fits are shown in Figure S3. ℒ and AICc give the log likelihood score and the correlated Akaike information criterion value, respectively. Best ℒ (maximum) and AICc (minimum) scores are bolded in the table. High (perhaps unrealistic) mutation rates are highlighted in italic.

